# Host-microbiome protein-protein interactions reveal mechanisms in human disease

**DOI:** 10.1101/821926

**Authors:** Hao Zhou, Juan Felipe Beltrán, Ilana Lauren Brito

## Abstract

Host-microbe interactions are crucial for normal physiological and immune system development and are implicated in a wide variety of diseases, including inflammatory bowel disease (IBD), colorectal cancer (CRC), obesity, and type 2 diabetes (T2D). Despite large-scale case-control studies aimed at identifying microbial taxa or specific genes involved in pathogeneses, the mechanisms linking them to disease have thus far remained elusive. To identify potential mechanisms through which human-associated bacteria impact host health, we leveraged publicly-available interspecies protein-protein interaction (PPI) data to find clusters of microbiome-derived proteins with high sequence identity to known human protein interactors. We observe differential targeting of putative human-interacting bacterial genes in metagenomic case-control microbiome studies. In nine independent case studies, we find evidence that the microbiome broadly targets human proteins involved in immune, oncogenic, apoptotic, and endocrine signaling pathways in relation to IBD, CRC, obesity and T2D diagnoses. This host-centric analysis strategy provides a mechanistic hypothesis-generating platform for any metagenomics cohort study and extensively adds human functional annotation to commensal bacterial proteins.

**One-sentence summary:** Microbiome-derived proteins are linked to disease-associated human pathways by metagenomic and protein-protein interaction analyses.

## Main Text

Metagenomic case-control studies of the human gut microbiome have implicated bacterial genes in a myriad of diseases. Yet, the sheer diversity of genes within the microbiome (Li et al., 2014) and the limitations of functional annotations (Joice et al., 2014) have thwarted efforts to identify the mechanisms by which bacterial genes impact host health. In the cases where functional annotations exist, they tend to reflect the proteins’ most granular molecular functions (*e.g.* DNA binding, post-translational modification) rather than their role in biological pathways (Lloyd-Price et al., 2017) and few, if any, relate to host cell signaling and homeostasis. Associating any commensal bacterial gene and a host pathway has thus far required experimental approaches catered to each gene or gene function (Nešić et al., 2014; Plovier et al., 2017).

Protein-protein interactions (PPIs) have revealed the mechanisms by which pathogens interact with host tissue through in-depth structural studies of individual proteins (Guven-Maiorov et al., 2017a; Hamiaux et al., 2006; Nešić et al., 2014), as well as large-scale whole-organism interaction screens (Dyer et al., 2010; Shah et al., 2018). These interactions are not limited to pathogens as many canonical protein-mediated microbe-associated molecular patterns (MAMPs) that directly trigger host-signaling pathways through pattern recognition receptors present on epithelial and immune tissues (Bhavsar et al., 2007) are conserved between pathogens and commensals (Lebeer et al., 2010), such as that between flagellin with Toll-like receptor 5 (TLR5). There is a growing recognition of the role for commensal-host PPIs in health (Table 1, Table S1): the *Akkermansia muciniphila* protein p9 binds intercellular adhesion molecule 2 (ICAM2) to increase thermogenesis and glucagon-like peptide-1 (GLP-1) secretion, a therapeutic target for type 2 diabetes (T2D) (LeValley et al., 2020); the protein Fap2 from *Fusobacterium nucleatum* binds T cell immunoreceptor with Ig and ITIM domains (TIGIT), inhibiting natural killer cytotoxicity; and a slew of ubiquitin mimics encoded by both pathogens (Guven-Maiorov et al., 2017b) and gut commensals (Stewart et al., 2018) play a role in modulating membrane trafficking. Whereas these efforts have progressed on a one-by-one basis, we hypothesized that host-microbiome PPIs that underlie health status may be widespread and that a systems-level approach could serve to provide additional information, through annotation of human pathways, about the role of bacteria in modulating health.

**Table 1.**
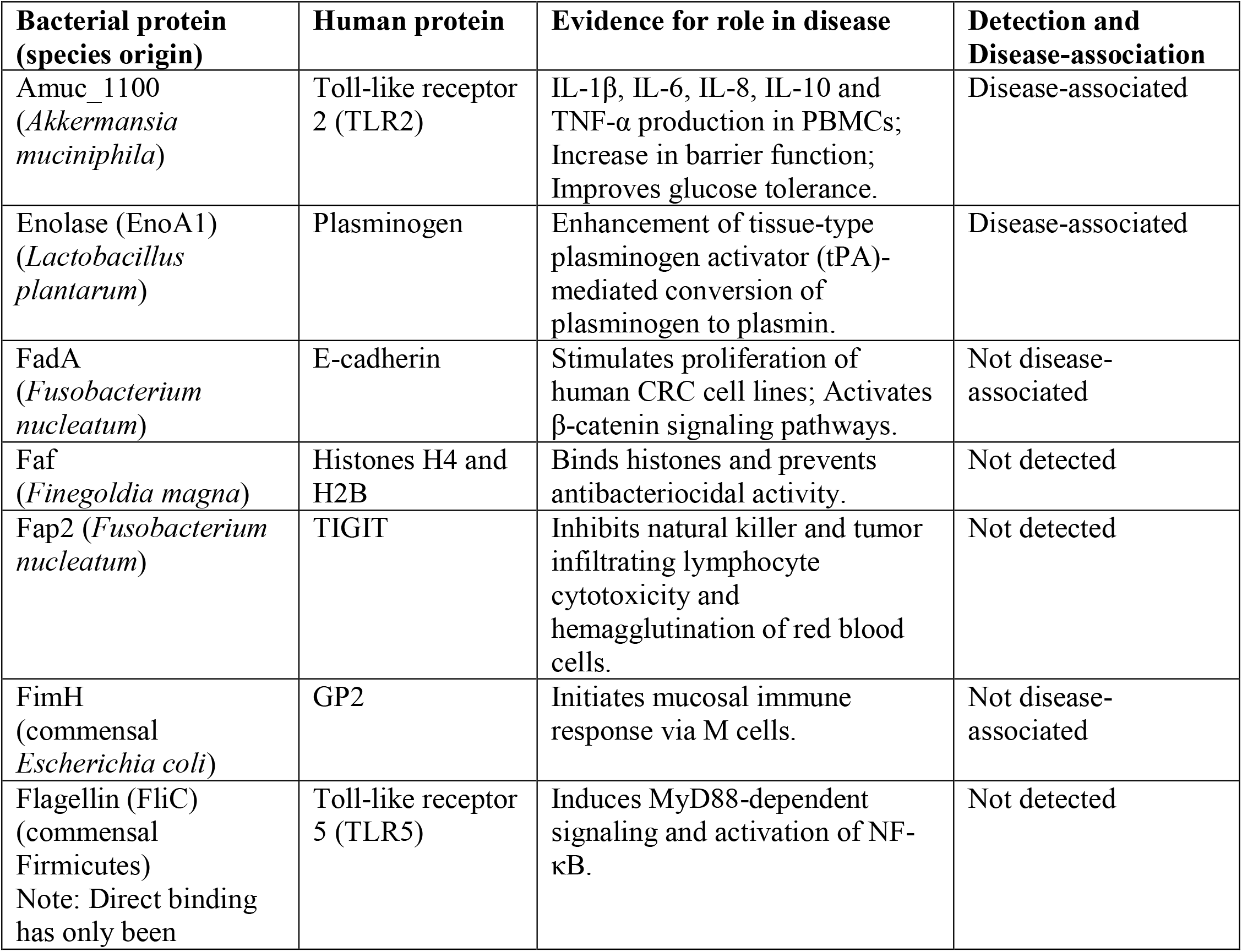
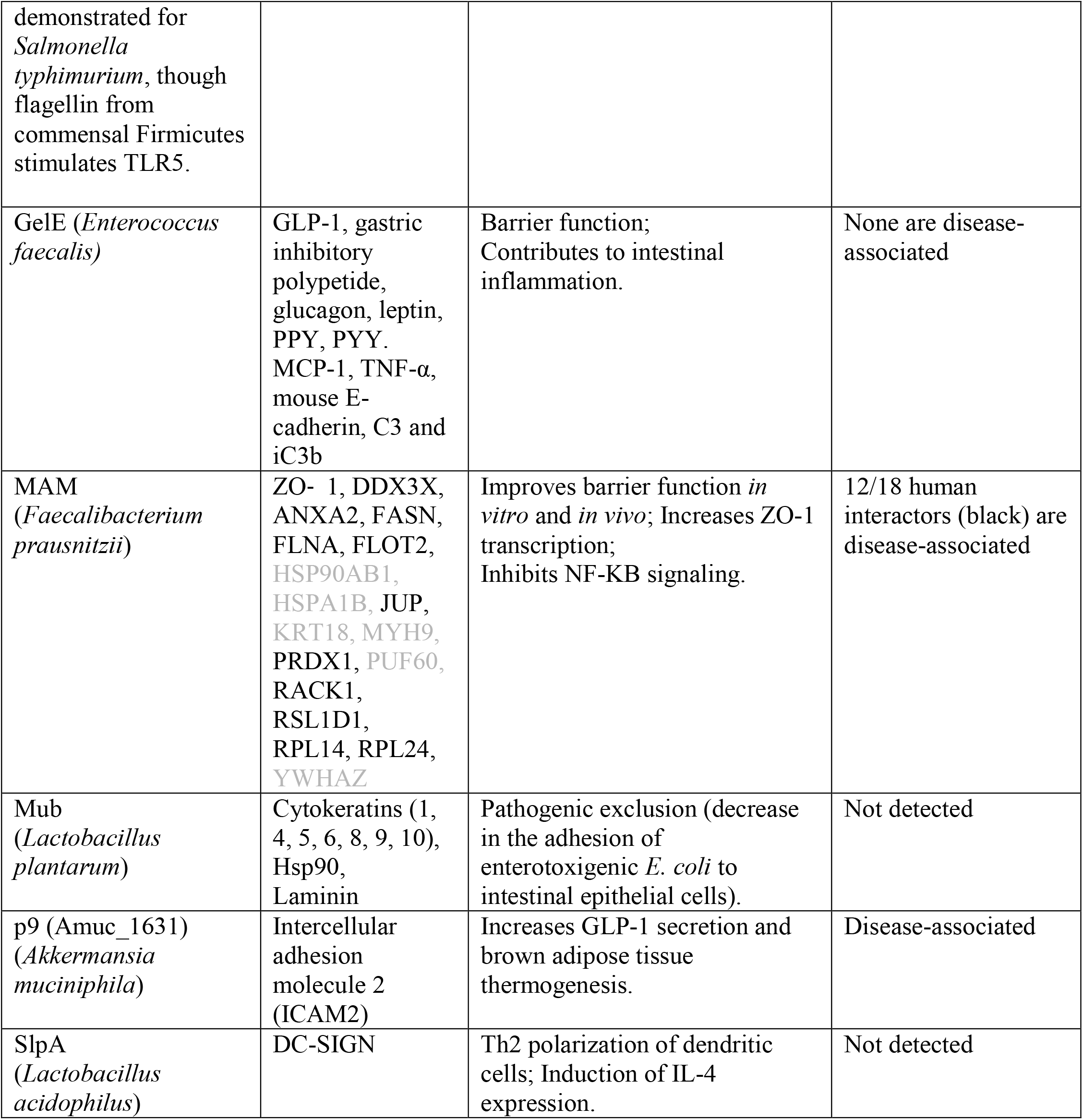
Examples of experimentally-verified host-microbiome PPIs that affect human cellular physiology and/or health. Designations include whether the bacterial proteins were detected within the nine metagenomic studies included in this analysis, and, if so, whether the human proteins were identified by our method as ‘disease-associated’. Extended information and citations are provided in Table S1.

Currently, few experimentally-verified inter-species PPIs exist involving human proteins, totaling 15,252 unique interactions in IntAct (Orchard et al., 2014), BioGRID (Oughtred et al., 2019), HPIdb (Ammari et al., 2016) and a set of manually curated PPI datasets (Fig. S1). Only a handful of these involve proteins pulled from the human gut microbiome. Expanding the commensal-human interaction network through state-of-the-art structural modeling (Guven-Maiorov et al., 2019) is untenable, as there are few sequences homologous to genes found in metagenomes represented in cocrystals from the Protein Data Bank (Burley et al., 2017) (PDB) (Fig. S2). In the absence of structure and experimental data, sequence identity methods have been used to great effect to infer host-pathogen PPI networks for single pathogens (Eid et al., 2016; Huo et al., 2015; Sen et al., 2016), but such approaches have not yet been applied at the community-level, as would be required for the human gut microbiome. Concerned over the reliability of interactions, we posited that we could leverage metagenomic case-control studies to hone in on those interactions relevant to disease, by focusing only on those interactions relevant to disease by virtue of their putative interactions with human proteins.

### Mapping microbiome proteins to known PPIs identifies potential mechanistic links to disease

All pathogen-host interactions are initially implicated in virulence, whereas microbiome-associated disorders tend not to follow Koch’s postulates (Byrd and Segre, 2016). To distinguish PPIs that may be associated with health versus disease, we compared community-level PPI profiles in large case-control cohorts of well-established microbiome-associated disorders—namely inflammatory bowel disease (IBD) (Franzosa et al., 2019; Schirmer et al., 2018), colorectal cancer (CRC) (Feng et al., 2015; Hannigan et al., 2018; Yu et al., 2017; Zeller et al., 2014), obesity (Le Chatelier et al., 2013), and T2D (Karlsson et al., 2013; Qin et al., 2012) (Fig. 1A, Table S2). In order to build community-level PPI profiles, we associated gene family abundances in these nine studies to a newly constructed database of bacterial human-protein interactors and the bacterial members of their associated UniRef clusters (Fig. S1), which represent homeomorphic protein superfamilies through sequence identity (Wu et al., 2004). For further assurance, we required microbiome proteins to have high amino-acid similarity (at least 70%) with the specific proteins with experimental evidence of interacting with human proteins. We noticed that proteins present exclusively in pathogenic organisms, such as the *Clostridium difficile* toxin B (TcdB) which binds frizzled 2 (FZD2), or expressed predominantly by pathogenic isolates, such as *Finegoldia magna* protein L, which binds immunoglobulin L chains, are consequently filtered out (Åkerström and Björck, 2009). We found that interspecies bacterial-human protein interface residues, in general, are highly similar, or even identical, between members of the same UniRef cluster filtered in the same manner (Fig. S3). Focusing on putative microbiome interactors with strong associations with disease weeds out a greater percent of interactions initially detected by yeast-2-hybrid (Y2H) methods and enriches for those that are based on affinity techniques (Fig. S4), and consequently removes the most “sticky” bacterial proteins (Fig. S5). The human protein with the highest degree remaining is nuclear factor NF-κB p105 subunit (NFKB1), a protein involved in immunodeficiency and bacterial infection, which was differentially targeted in CRC (in Vogtmann et al.). After applying a random forest classifier trained on each disease cohort (Fig. S6), we find 1,102 commensal bacterial protein clusters associated with disease, by virtue of their putative interactions with 648 human proteins (Table S3).

**Figure 1.**
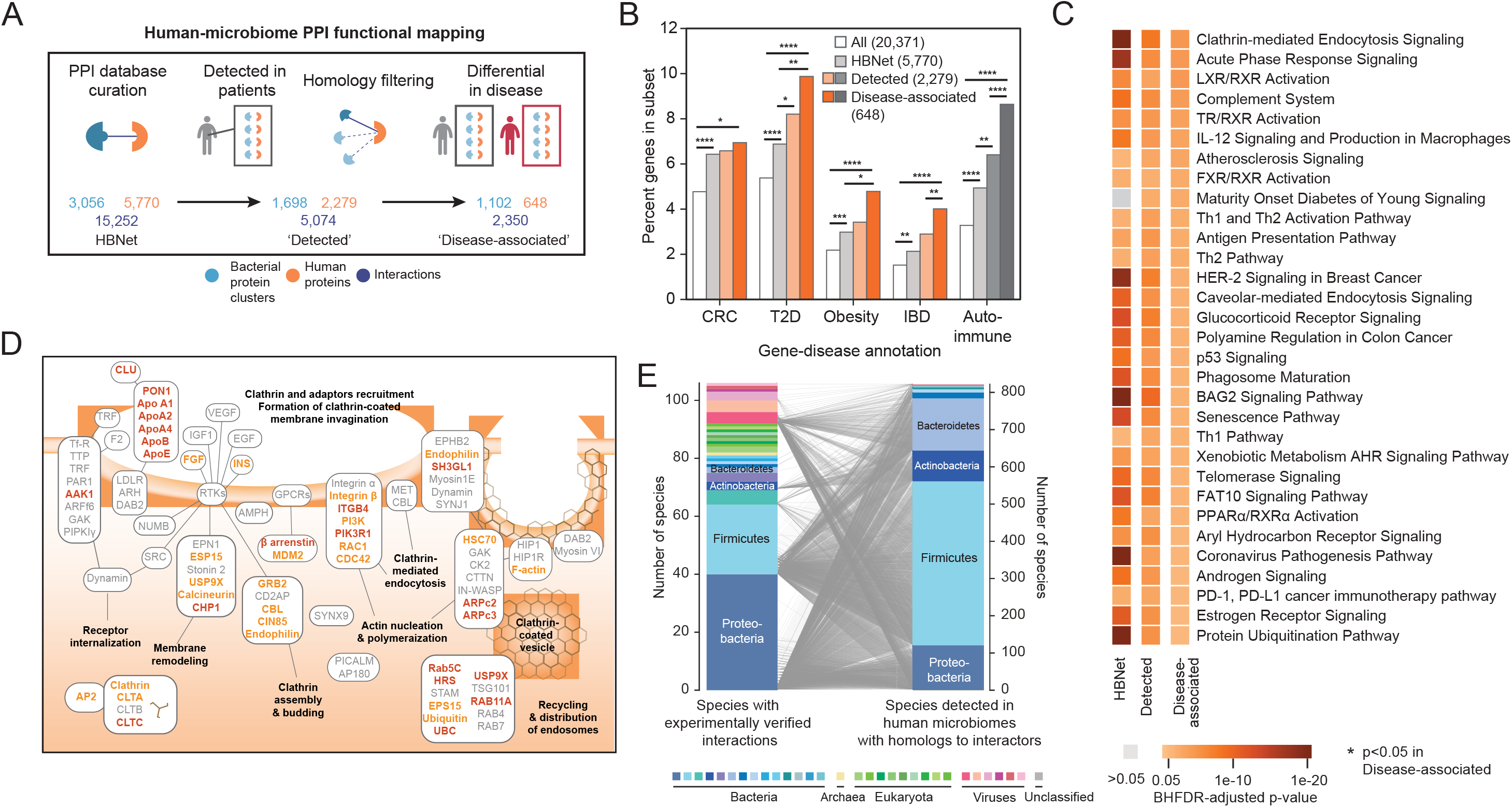
Human proteins differentially targeted by the microbiome in disease are enriched for relevant gene-disease associations. (A) The number of interspecies bacterial protein clusters (blue), human proteins (orange) and interactions (dark blue) in the human-bacteria PPI network; the number of bacterial protein clusters detected in patients from nine metagenomic studies that also have homology to experimentally-verified interactors and their putative human interactors; and the number of bacterial clusters and human proteins associated with disease through our metagenomic machine learning approach, by comparing abundances in cases (grey) and control (red). (B) Proportions of human proteins implicated in disease, according to their GDAs (GDAs > 0.1) in DisGeNET, within: all reviewed human proteins; HBNet; human interactors with detected bacterial proteins; and those human interactors with feature importances above the 90^th^ percentile in their respective cohorts. p-values for enrichments are depicted by: * p<0.05; ** p<0.01; *** p<10-3; **** p<10-4 (Chi-square test). Total numbers of each set are noted in the legend. (C) Human cellular pathways (annotated by IPA) enriched in the set of human proteins within HBNet (left) and those detected across all nine metagenomic case-control studies (right) colored according to their Benjamini-Hochberg false discovery rate (BHFDR)-adjusted p-value. Only those pathways with BHFDR-adjusted < 0.05 in the disease-associated sets are shown. p-values for enrichments are depicted by: * p<0.05; ** p<0.01; *** p<10-3; **** p<10-4 (Fisher’s Exact test). (D) All human proteins within the Clathrin-Mediated Endocytosis Signaling pathway, as annotated by IPA, are depicted. Protein targets detected in the nine metagenomic studies are highlighted in orange. Those in the Disease-associated subset are in brown. Specific interactions and the nature of interactions were simplified, with boxes roughly representing proteins within the same signaling cascade and/or complex. (E) 106 species (left) with experimentally verified proteins in 3,056 bacterial protein clusters are mapped to 821 bacterial species (right) with homologs detected in patients’ metagenomes (right), representing a total of 1,698 clusters. Species are colored according to phylum.

Surprisingly, within the human proteins associated with CRC via the microbiome are a number of previously identified CRC-associated genetic loci (*e.g.* immunoglobulin 8 (IL-8), toll-like receptor 2 (TLR2), selenoprotein P, the phospholipid scramblase 1, MDM4, and the histone acetyltransferase p300, among others. This represents a larger trend: moving from the 5,770 human proteins within the interaction network (‘HBNet’), to the 2,279 human proteins with bacterial interactors detected in human microbiomes (‘Detected’), to the 648 that are associated with disease (‘Disease-associated’), we observe increasing enrichment for proteins with previously-reported gene-disease associations (GDA) in CRC, diabetes, obesity, and IBD (Fig. 1B). These enrichments are even more pronounced when examining each specific disease cohort (Fig. S7). However, we see enrichment for microbiome-associated disorders in each of the cohorts, reflecting their associated relative risks (Jess et al., 2019; Jurjus et al., 2016; Kang et al., 2019; de Kort et al., 2017; Stidham and Higgins, 2018). In fact, out of all of the proteins with any GDA in the disease-associated set, 45.2% percent have more than one GDA for our diseases of interest. We suspected this may extend to autoimmune diseases, which are increasingly studied in the context of the gut microbiome (Gianchecchi and Fierabracci, 2019), and we confirm enrichment of GDAs for autoimmune disorders in the human proteins implicated by our method (Fig. 1B, Fig. S7). This concordance between known disease annotation and disease association demonstrates the utility of using PPIs to capture molecular heterogeneity that underlies microbiome-related disease.

In evaluating the statistical significance of recurrent human functional annotations, we performed pathway enrichment analysis on the implicated human proteins and find proteins with established roles in cellular pathways coherent with the pathophysiology of IBD, CRC, obesity and T2D (Fig. 1C), namely those involving immune system, apoptosis, oncogenesis, and endocrine signaling pathways. Most enriched pathways include human proteins across the four types of disease cohorts analyzed, reflecting their associated relative risks (Jess et al., 2019; Jurjus et al., 2016; Kang et al., 2019; de Kort et al., 2017; Stidham and Higgins, 2018). Human proteins differentially targeted by microbiome-sourced proteins have roles in pathways involved in bacterial pathogenesis and underlying inflammation, such as the IL-12 signaling pathway and clathrin-mediated endocytosis signaling. These pathways were expected due to shared evolutionary histories between the screened pathogens and gut microbiota and opportunism within the microbiome. The involvement in the clathrin-mediated endocytosis pathways (Fig. 1D) further hints at how commensal proteins may enter human cells. Pathways related to bile salt metabolism and cholesterol metabolism (LXR/RXR, TX/RXR and FXR/RXR activation pathways), which are also tied to immune evasion (Alatshan and Benkő, 2021; Valledor et al., 2004) are also enriched, expanding the role of the microbiota in these pathways beyond their enzymatic functions.

Within these pathways, we see specific examples of known molecular mechanisms for these diseases now implicated with microbiome-host PPIs: Actin-related protein 2/3 complex subunit 2 (ARPc2) (associated in the Schirmer et al., Feng et al., Yu et al. and Zeller et al. cohorts) regulates the remodeling of epithelial adherens junctions, a common pathway disrupted in IBD (Franke et al., 2008). We see the targeting of mitogen-activated protein kinase kinase kinase kinase 1 (MAP4K1) enriched in the Zeller *et al.* CRC cohort, which is in line with its role in inflammation (Chuang et al., 2016). DNA methyltransferase 3a (DNMT3A) is involved in chromatin remodeling and has been shown to be important for intestinal tumorigenesis (Weis et al., 2015), serve as a risk loci in genome-wide association studies (GWAS) studies for Crohn’s disease (Franke et al., 2010), mediates insulin resistance (You et al.) and has aberrant expression in adipose tissue in mice (Kamei et al., 2010). Concordantly, it was associated with the CRC, IBD, T2D and obesity microbiome studies we examined (Feng et al., Yu et al., Zeller et al., LeChatelier et al., Qin et al. and Schirmer et al.). This host-centric annotation is useful beyond large-scale analysis of metagenomic data, as it broadly enables hypothesis-driven research into the protein-mediated mechanisms underlying microbiome impacts on host health.

Although the set of experimentally-verified interactions (HBNet) includes interactions originating from 82 unique bacterial species, an initial concern was that a disproportionate number of bacteria-human PPIs are derived from high-throughput screens performed on a smaller number of intracellular pathogens, *e.g*. *Salmonella enterica* (Walch et al., 2021), *Yersinia pestis* (Dyer et al., 2010; Yang et al., 2011), *Francisella tularensis* (Dyer et al., 2010)*, Acinetobacter baumannii* (Schweppe et al., 2015), *Mycobacterium tuberculosis* (Penn et al., 2018), *Coxiella burnetii* (Wallqvist et al., 2017), *Chlamydia trachomatis* (Mirrashidi et al., 2015) and *Legionella pneumophila* (Yu et al., 2015), *Burkholderia mallei* (Memisević et al., 2013), and *Bacillus anthracis* (Dyer et al., 2010); as well as one extracellular pathogen *Streptococcus pyogenes* (Happonen et al., 2019) (Table S4). Despite this bias, we find that homologs detected in patient microbiomes come from a set of 821 species that better reflects the phyla typically associated with human gut microbiomes (Fig. 1E).

### Microbiome proteins access human proteins by various means

We next examined the localization of human protein targets. Amongst those human proteins in the detected and disease-associated sets, we saw increasing enrichment of genes expressed in epithelium, liver, adipose tissue and blood components (Fig. 2A). Although we presume many of the interactions occur within in the epithelial layer of the gastrointestinal tract, disease-associated human interactors were not especially localized to gastrointestinal tissue, nor any tissue in particular, with the exception of bone marrow (p=0.047, chi-square test) (Fig. S8). Impaired intestinal barrier function and the translocation of commensal bacteria, both of which feature in the pathogenesis of IBD (Ahmad et al., 2017), CRC (Genua et al., 2021) and other microbiome-associated disorders (Ruff et al., 2020), allow bacterial proteins to access tissues exterior to the gut. Nevertheless, we suspect that the absence of enrichment in gut tissues largely reflects the human tissues, cells, and fluids used for experimental interaction screening (*e.g*. HeLa cells (Walch et al., 2021), HEK293T (Mirrashidi et al., 2015), macrophages (Walch et al., 2021), plasma (Happonen et al., 2019), saliva (Happonen et al., 2019), spleen (Dyer et al., 2010; Yang et al., 2011), and lung (Schweppe et al., 2015)), thereby selecting proteins with more general expression patterns. This data underscores the need for screening using gastroenterological protein libraries to identify gut-specific host-microbiome PPIs.

**Figure 2.**
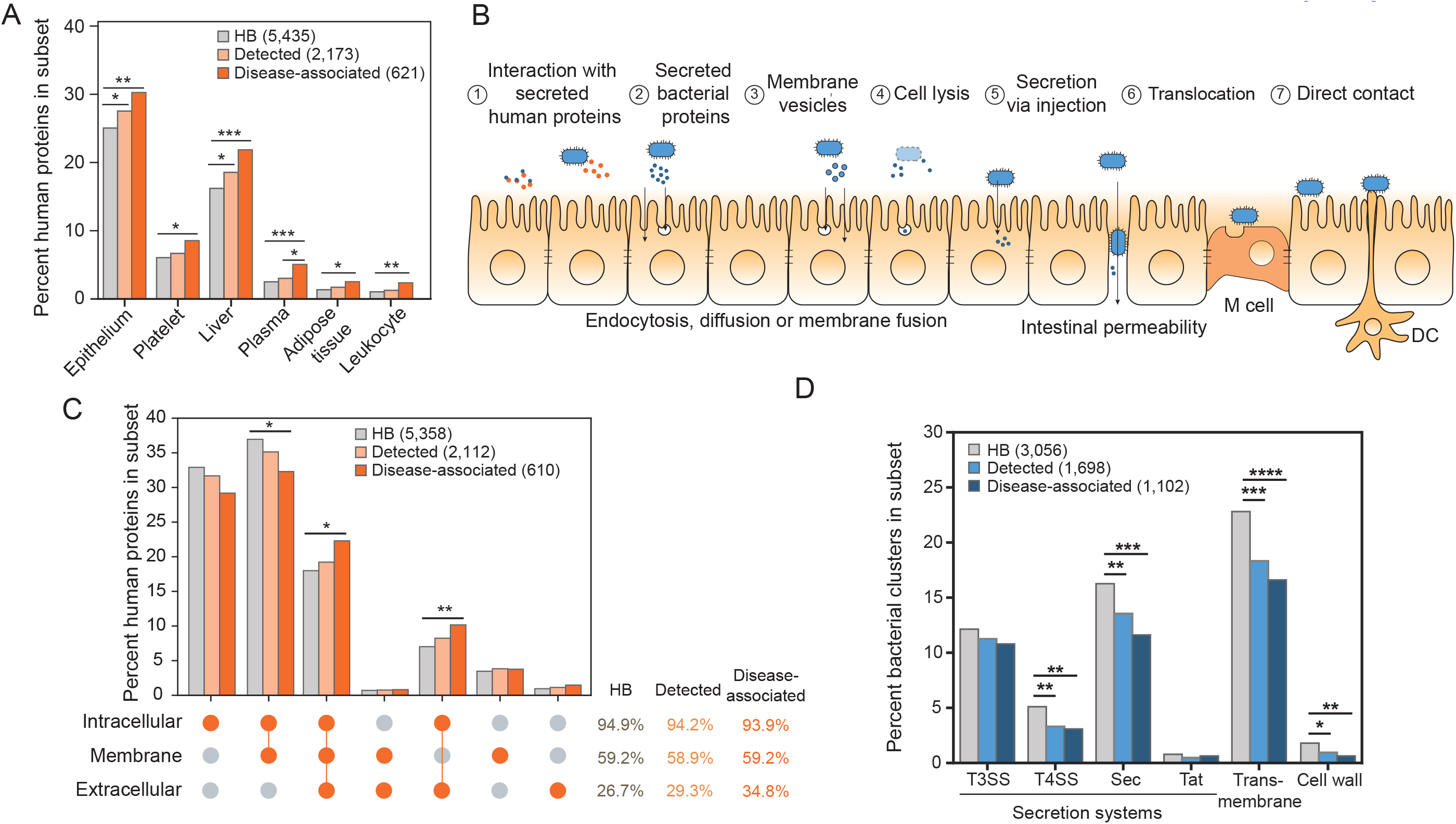
Bacterial proteins gain access to human proteins through a variety of mechanisms. (A) Proportions of human proteins in the HBNet, Detected and Disease-associated subsets are plotted according to their enrichments in tissues and fluids, as annotated using DAVID. Only those with significant enrichment between any two subsets are shown. p-values for enrichments are depicted by: * p<0.05; ** p<0.01; *** p<0.001; **** p<0.0001 (EASE Score provided by DAVID, a modified Fisher Exact P-value; FDR-adjusted). Total numbers of each set are noted in the legend. (B) A schematic depicting potential opportunities for bacterial proteins to access human proteins. Interactions may involve: (1) secreted human proteins, (2) bacterial proteins secreted into the extracellular space; (3) membrane vesicles that are endocytosed or can fuse with human cell membranes; (4) bacterial cellular lysate; (5) proteins injected into human cells by T3SS, T4SS and T6SS, (6) cells and their products that translocate as a result of barrier dysfunction or “leaky gut”, and/or (7) direct contact with M cells, dendritic cells (DC), or epithelial cells. (C) Proportions of human proteins in the HBNet, Detected and Disease-associated subsets, are plotted according to their subcellular locations, as annotated using Gene Ontology Cellular Component, is depicted. p-values for enrichments are depicted by: * p<0.05; ** p<0.01; *** p<0.001; **** p<0.0001 (Chi-square test). Total percentages for these subsets is listed at right, along with p-values. Total numbers of each set are noted in the legend. (D) Proportions of bacterial gene clusters in the HBNet, Detected and Disease-associated subsets are plotted according to their transmembrane and secretion predictions, annotated using TMHMM, EffectiveDB and SignalP. p-values for enrichments are depicted by: * p<0.05; ** p<0.01; *** p<0.001; **** p<0.0001 (Chi-square test). Total numbers of each set are noted in the legend.

At the cellular level, microbial proteins can access human proteins via several well-established means (Fig. 2B). Canonical MAMPs tend to involve surface receptors (*e.g*. TLRs, Nod-like receptors), which comprise 59.2% of the disease-associated interactors (Fig. 2C), although we cannot confirm their orientation. We expect that this may be an underestimate of the interactions involving human membrane interactors, as solubility issues preclude their representation in interaction screens. In addition to canonical MAMP receptors, newly described surface receptors include: adhesion G protein-coupled receptor E1 (ADGRE1), a protein involved in regulatory T cell development (Lin et al., 2005); and receptor-type tyrosine-protein phosphatase mu (PTPRM), involved in cadherin-related cell adhesion (Brady-Kalnay et al., 1995), among others. Alternatively, several established host-microbiome PPIs (Table 1) involve human proteins that are secreted, such as the extracellular matrix protein laminin (Singh et al., 2018) and immune modulators, such as extra-cellular histones (Brinkmann et al., 2004; Murphy et al., 2014). Secreted proteins make up 34.8% of the disease-targeted human interactors, and include these, in addition to the cytokine IL-8, galectin-3, and complement 4A.

Interestingly, a large number of disease-associated human interactors (178 proteins, or 29.1%) are exclusively intracellular (Fig. 2C), suggesting additional interaction schemes. MAM (microbial anti-inflammatory molecule), a secreted protein from *Faecalibacterium prausnitzii,* can inhibit NF-κB signaling and increase tight junction integrity, whether it is introduced via gavage in mouse models, or when it is ectopically expressed from within intestinal epithelial cells *in vitro* (Xu et al., 2020), suggesting that it is uptaken by cells *in vivo*. Bacterial products or, in some cases, intact bacteria, may be endo-, pino- or transcytosed, a process that can be initiated by receptors (Malyukova et al., 2009; Tan et al., 2015), allowing bacterial proteins to access cytoplasmic and even nuclear targets. Alternatively, membrane vesicles, decorated with proteins and carrying periplasmic, cytoplasmic and intracellular membrane proteins as cargo, can be uptaken by human cells via endocytosis or membrane diffusion (Jones et al., 2020). Although membrane vesicles have been well-documented in Gram-negative bacteria, an example of vesicle production by Gram positive segmented filamentous bacteria was recently shown to interact with intestinal epithelial cells and promote the induction of Th17 cells (Ladinsky et al., 2019).

Accordingly, bacterial proteins interacting with human secreted and surface proteins would be expected to contain signatures of surface localization or extracellular secretion. Indeed, we find that 12.2% of the disease-associated microbiome proteins are predicted to contain signal peptides allowing for secretion by the Sec or Tat pathways (Fig. 2D), which are ubiquitous across phyla (Fig. S9). These systems typically work alongside additional secretion systems to situate proteins in the cell membrane or secrete them extracellularly, though their associated signal peptides are more difficult to predict (Green and Mecsas, 2016; Hui et al., 2021). Another 16.6% of disease-associated proteins are predicted to be transmembrane, albeit with unknown orientation, potentially allowing for direct contact with live or intact bacteria, or bacterially-produced membrane vesicles. A small number of proteins were found destined for the cell wall (Fig. 2D). To our surprise, secreted and surface proteins were found to be negatively enriched in the disease-associated bacterial interactors.

Finally, type 3, type 4 and type 6 secretion systems (T3SS, T4SS and T6SS) can be used to secrete proteins directly into human cells. Proteins with T3SS and T4SS signals make up a significant (13.6%), albeit diminishing portion of the disease-associated microbiome proteins (Fig. 2D). These proteins are mostly derived from gut Proteobacteria, to which these systems are generally restricted (Abby et al., 2016) (Fig.2D, Fig. S9). Based on the bacterial cluster representatives from in the microbiomes from these nine cohorts, we find evidence that at least 79.0% and 58.9% of disease-associated clusters predicted to be secreted by T3SS and T4SS, respectively, have representative proteins found in organisms with the corresponding secretion systems (T6SS were excluded due to the limited availability of prediction tools). Nevertheless, the extent to which these systems, and orthologous systems in Gram positive bacteria (Madden et al., 2001), play a role in host-microbiome protein trafficking remains unknown. In total, this data suggests that there is not one single mechanism dominating host-microbiome interactions, but that interactions are facilitated by several means.

### Microbiome proteins gain host-relevant “moonlighting” annotations

One of the major advantages of our work is that through this new interaction network, we vastly improve our ability to annotate host-relevant microbiome functions. 13.5% of our disease-associated bacterial clusters contain no members with annotated microbial pathways/functions in KEGG (Kyoto Encyclopedia of Genes and Genomes) (Kanehisa et al., 2017) (Fig. 3A). Using similar homology searching against bacterial interactors, most of these genes can now be annotated according to the pathways of their human targets, obtaining a putative disease-relevant molecular mechanism (Fig. S10). Interestingly, most of the bacterial clusters with KEGG pathway annotations also gain a secondary human pathway annotation. Of those that could be annotated, disease-associated clusters are involved primarily in translation and central metabolism (Fig. 3B). This dual function is not entirely surprising, as a number of these have orthologs that have been previously identified as bacterial ‘moonlighting’ proteins, which perform secondary functions in addition to their primary role in the cell (Henderson, 2014). *Mycoplasma pneumoniae* GroEL and *Streptococcus suis* enolase, a protein involved in glycolysis, bind to both human plasminogen and extra-cellular matrix components (Hagemann et al., 2017; Henderson and Martin, 2013). *Mycobacterium tuberculosis* DnaK signals to leukocytes causing the release of the chemokines CCL3-5 (Lehner et al., 2000). *Streptococcus pyogenes* glyceraldehyde-3-phosphate dehydrogenase (GAPDH), canonically involved in glycolysis, can be shuffled to the cell surface where it plays a role as an adhesin, and can also contribute to human cellular apoptosis (Seidler and Seidler, 2013). These examples distinctly illustrate how bacterial housekeeping proteins are used by pathogens to modulate human health. In this study, we uncover commensal proteins that similarly may have ‘interspecies moonlighting’ functions and appear to be pervasive throughout our indigenous microbiota.

**Figure 3.**
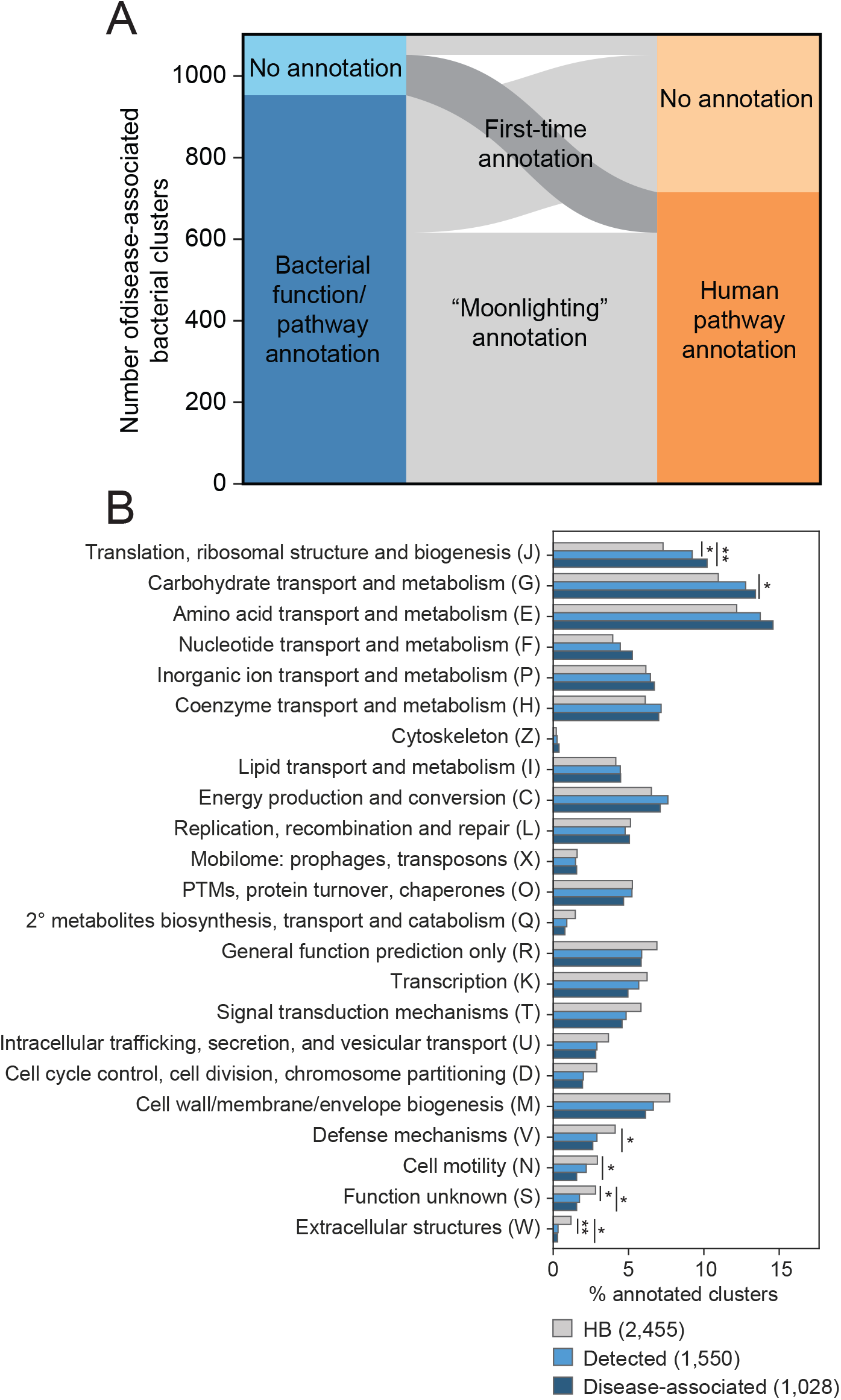
Human pathway annotation can be propagated through interactors to improve bacterial pathway annotation. (A) Paired stacked bar plots showing the 1,102 disease-associated bacterial protein clusters according to whether they are able to be annotated by KEGG (left) and their inferred pathways according to the human proteins they target (right), as annotated by WikiPathways (Slenter et al., 2018). (B) Proportions of the bacterial clusters in the HBNet, Detected and Disease-associated subsets according to their COG functional categories are plotted. p-values are depicted by: * p<0.05; ** p<0.01; *** p<0.001; **** p<0.0001 (Chi-square test). Total numbers of each set are noted in the legend.

### Microbiome proteins may act on human targets as therapeutic drugs

There is direct evidence for at least two commensal proteins which induce physiological effects on the host when delivered by oral gavage: purified *A. muciniphila* Amuc_1100 and *F. prausnitzii* MAM to ameliorate glucose intolerance and colitis, respectively (Plovier et al., 2017; Xu et al., 2020). We suspect that this may extend to additional commensal proteins. Consistent with this idea, we find that indeed many disease-associated human proteins are known drug targets (Table S5). For example, nafamostat mesylate is an anticoagulant that can bind complement protein C1R, suppresses coagulation and fibrinolysis and provides protection against IBD (Isozaki et al., 2006) and CRC (Lu et al., 2016). These human proteins are also differentially targeted in healthy patients by the transcriptional regulator spo0A in Lactobacilli, Streptococci and *F. prausnitzii* (Fig. 4A, Table S6). Imatinib mesylate (brand name: Gleevec) targets several Src family tyrosine kinases, including LCK, which is involved in T cell development and has a recognized role in inflammation (Kumar Singh et al., 2018). Bacterial proteins targeting these same kinases are consistently enriched in healthy controls across both IBD and three CRC cohorts we analyzed (Fig. 4B, Table S6). In addition, imatinib can also halt the proliferation of colonic tumor cells and is involved generally in inflammatory pathways, through its inhibition of TNF-alpha production (Wolf et al., 2005).

**Figure 4.**
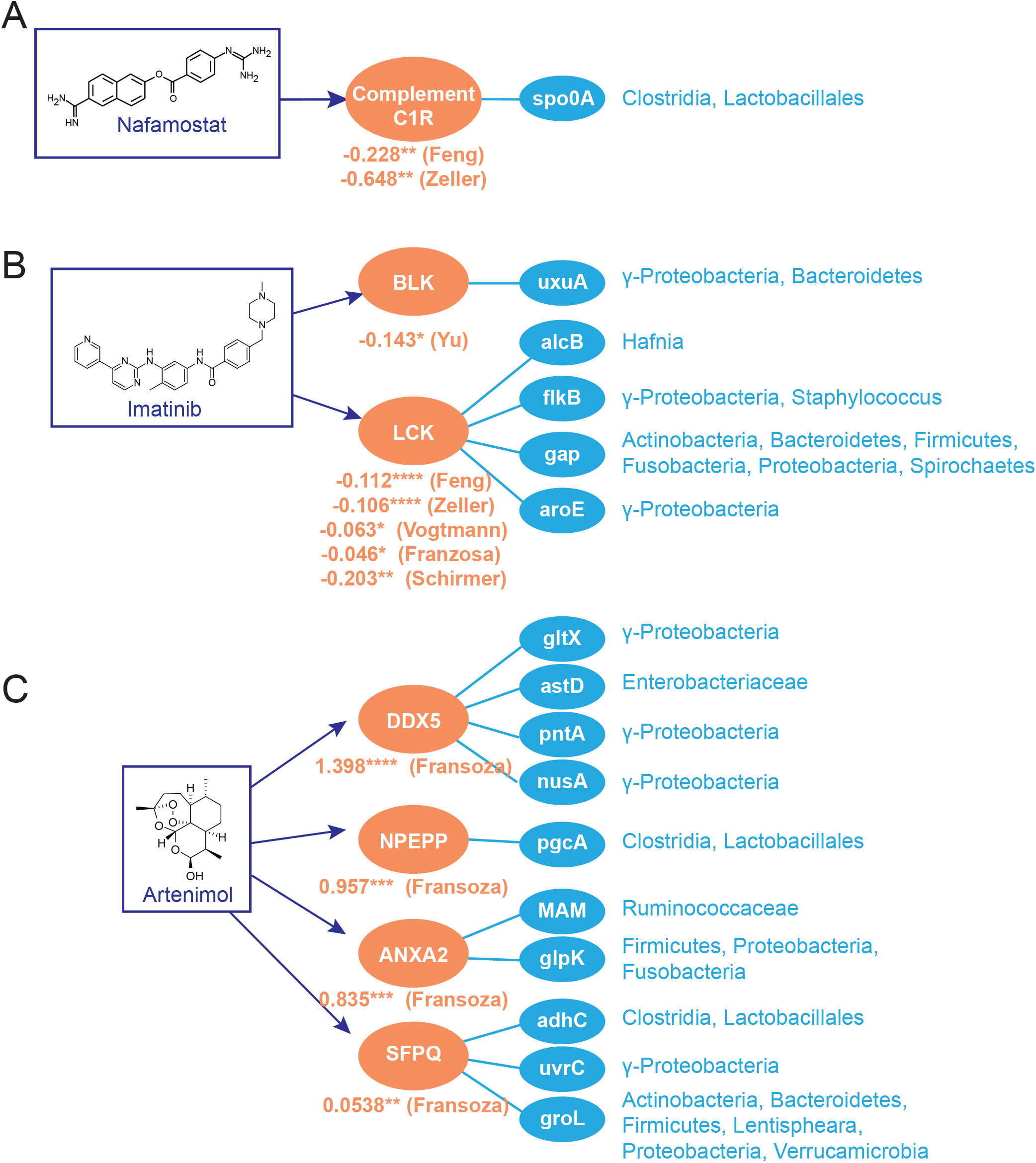
Human proteins targeted by gut commensal proteins include known therapeutic drug targets. (A) Nafamostat, (B) imatinib and (C) artenimol target human proteins that are differentially targeted by bacterial proteins detected in the stated metagenomic studies. Log10 relative mean summed abundances of bacterial interactors in patients versus controls are provided. p-values were calculated by the Mann-Whitney rank-sum test, * p<0.05; ** p<0.01; *** p<10^−3^; **** p<10^−4^). Full taxa and UniRef numbers for all bacterial proteins shown are provided in Table S6.

We also find instances where the off-label effects or side effects associated with the drug match our microbiome-driven human protein association. For instance, the antimalarial drug artinemol targets human proteins that were found to be differentially targeted by IBD cohorts’ microbiomes (in Franzosa *et al.*): the RNA helicase DDX5, puromycin-sensitive aminopeptide (NPEPP), annexin A2 (ANXA2) and the splicing factor SFPQ (Fig.4C, Table S6). Whereas artinemol and related analogs have been shown to be effective at preventing dextran sulfate-induced colitis in mice (Hu et al., 2014; Yan et al., 2018) and wormwood, its natural source, has been established as a herbal treatment for IBD (Krebs et al., 2010), microbiota-derived proteins have greater association with IBD patients, suggesting that artinemol and commensal proteins may be acting on the same targets in opposing ways. Whereas the notion of microbiome-derived metabolites acting as drugs is well-appreciated (Donia and Fischbach, 2015; Wilson et al., 2019), this work broadens the scope of microbiome-derived drugs to include protein products acting through PPI.

## Discussion

Here, we reveal an extensive host-microbiome PPI landscape. To achieve this, we benefit from existing methods in pathogen-host PPI discovery, further informed by community-level PPI profiles of genes differentially detected in human metagenomes. This work highlights host mechanisms targeted by the gut microbiome and the extent to which these mechanisms are targeted across microbiome-related disorders. However, this network is far from complete. Few of the studies on which this interaction network is based were performed on commensal bacteria and intestinal tissue, and therefore, we may be missing interactions specific to our most intimately associated bacteria. In support of our method, among those host-microbiome PPIs that have been well-studied for both binding and their effect on human cellular physiology or disease pathophysiologies (Table 1, Table S1), we were able to associate over half of the PPIs with one or more metagenomic studies. In addition to large-scale PPI studies involving commensal bacteria and their hosts, further in-depth studies will be needed to fully characterize these mechanisms, such as whether these bacterial proteins activate or inhibit their human protein interactors’ pathways, and under what conditions these interactions take place.

This platform enables a high-throughput glimpse into the mechanisms by which microbes impact host tissue, allowing for mechanistic inference and hypothesis generation from any metagenomic dataset. Pinpointing microbe-derived proteins like this that interact directly with human proteins will enable the discovery of novel diagnostics and therapeutics for microbiome-driven diseases, more nuanced definitions of the host-relevant functional differences between bacterial strains, and a deeper understanding of the co-evolution of humans and other organisms with their commensal microbiota.

## Acknowledgments

We wish to acknowledge members of the Brito lab, Indrayudh Ghosal, Giles Hooker and Andy Clark for their thoughtful comments. Funding: Ilana Brito is a Pew Scholar in Biomedical Sciences, a Packard Foundation Fellow, and a Sloan Foundation Research Fellow. Ilana Brito is funded by the National Institutes of Health (1DP2HL141007).

## Author contributions

H.Z., J.F.B and I.L.B. conceptualized and designed the study and co-wrote the manuscript.

## Data and materials availability

All data used for this paper is publicly available and described in the Methods and in Table S2. Disease-associated human-microbiome PPIs are listed in Table S3.

## Methods

### Building a putative bacteria-human protein-protein interaction (PPI) network

Interactions were downloaded from the IntAct database (Orchard et al., 2014), HPIdb 3.0 (Ammari et al., 2016) and BioGRID (Oughtred et al., 2019) [June 2021], and supplemented with additional host-microbe interaction studies, whose interactions were added manually (PMIDs: 31227708, 34237247, 22213674, 18937849, 8900134, 17709412, 19047644, 23954158, 24335013, 24936355, 25680274, 26548613, 28281568, 29748286, 30072965, 30242281, 32566649, 32736072, 18808384, 22344444, 33820962, 31611645, 32051237, 18941224, 19627615, 3125250, 19752232, 21441512, 19542010, 11113124, 29335257, 21740499, 18541478, 9466265, 24204276, 23800426, 27302108, 25739981, 19907495, 31503404, 25118235, 25788290, 21699778, 26755725, 14625549). Only interactions with evidence codes that indicated binary, experimental determination of the interaction between UniProt identifiers with non-matching taxa were preserved, thereby excluding co-complex associations, small molecule interactions, and predicted interactions. Uniref100/90 clusters containing human proteins and Uniref50 cluster containing bacterial proteins were downloaded from UniProt [June 2021], to which interspecies protein interactors were mapped (Suzek et al., 2015). PPIs comprising one Uniref100/90 cluster containing human proteins and one Uniref50 cluster containing bacterial proteins were retained for downstream analyses. Within each UniRef50 bacterial cluster, we further filtered the sequences such that only bacterial members of the cluster within 70% sequence similarity to the experimentally verified protein were labeled as putative interactors. Sequence similarity was calculated using a Smith-Waterman local alignment with the BLOSUM62 matrix via python’s parasail (Daily, 2016) library (v.1.1.17) and tallying the number of matches in the pairwise alignment that represent frequent substitutions (non-negative BLOSUM62 scores), divided by the length of the experimentally-verified interactor.

### Processing of metagenomic shotgun sequencing data

The datasets used in this study, with the exception of the PRISM dataset (Franzosa et al., 2019), were curated as part of ExperimentHub (Pasolli et al., 2017) (Table S1). Within each study, we removed samples that had abnormally low (less than 10^7^) reads. We downloaded all protein abundance matrices, annotated at the level of UniRef90 clusters via HUMAnN3 (Beghini et al., 2021), and associated metadata. For PRISM, we processed data in a parallel manner, as outlined in Pasolli et al. (Pasolli et al., 2017). For each study, we mapped UniRef90 bacterial clusters to UniRef50 clusters using DIAMOND (Buchfink et al., 2015) blastp, requiring greater than 90% sequence identity and greater than 90% coverage.

### Prioritization of disease-associated human targets

For each patient, we generate a file of human proteins representing the cumulative abundances of their putative bacterial protein interactors. In each study, we filtered out proteins present in fewer than 5% of the cohort. To identify host-microbiome interactions that associate with disease, processed abundance matrices of putative human interactors were used to train a random forest machine learning classifier on the task of separating case and control patients and, after verifying that they achieve reasonable performance on the task using five-fold cross-validation with grid search-based hyperparameters tuning for each study (Fig. S6), we extract the average feature importance from 100 iteratively trained class-balanced classifiers. Having used the scikit-learn (Pedregosa et al., 2011) implementation of the random forest algorithm, feature importance corresponds to the average Gini impurity of the feature in all splits that it was involved in. Gini feature importance is a powerful prioritization tool, as it can capture the multivariate feature importance (whereas simple metrics like log-odds ratio and corrected chi-squared statistics only capture univariate feature importance). We created a disease-associated set for the proteins that had feature importances above the top 90^th^ percentile. As an alternative to calculating human protein abundances by summing the total bacterial abundances of their interactors, we tested the effect of first normalizing bacterial abundances by their respective number of putative human interactors. This did not qualitatively change the conclusions drawn from our analyses.

### Human pathway annotation and enrichment analysis

Disease annotations were extracted from all of GDAs from DisGeNET (Piñero et al., 2017) (June 2021). We additionally downloaded all reviewed human proteins from Uniprot (Ding et al., 2018) (June 2021), annotating them in the same manner, in order to accurately compare background label frequencies. Lacking a simple hierarchy of disease, we binned similar disease terms into the 5 larger categories relevant to our study. Human protein identifier labels are provided in Supplementary Note 1. We performed pathway enrichment analysis using QIAGEN’s Ingenuity® Pathway Analysis software (IPA®, QIAGEN Redwood City, CA, USA, https://www.qiagen.com/ingenuity). Sets of human proteins (HBNET, Detected, Disease-associated) were uploaded as UniProt identifiers into the desktop interface and submitted to their webserver for Core Enrichment Analysis was conducted only on human tissue and cell lines and IPA’s stringent evidence filter. Pathways were considered enriched if they had Benjamini-Hochberg-corrected p values < 0.05. Subcellular locations for human proteins were obtained using GO Cellular Component terms associated with each protein in UniProt. We aggregated the following GO terms: Extracellular: Extracellular region (GO:0005576), Extracellular matrix (GO:0031012); Membrane: Cell surface (GO:0009986); Membrane (GO:0016020), Cell junction (GO:0030054); Cell projection (GO:0042995); and Intracellular: Cytoplasm (GO:0005737); Cell body (GO:0044297); Nucleoid (GO:0009295); Membrane-enclosed lumen (GO:0031974); Organelle (GO:0043226); Endomembrane system (GO:0012505); Midbody (GO:0030496). Tissue-specific RNA expression enrichment was performed using DAVID bioinformatics resources (Huang et al., 2009). Additionally, tissue-specific protein localization data was downloaded from Human Protein Atlas version 20.1 (Uhlen et al., 2010). We retained those with ‘enhanced’, ‘supported’ and ‘approved’ reliability. We additionally annotated all human proteins with any known drug targets from the DrugBank database (Wishart et al., 2018) and DrugCentral (June 2021) (Avram et al., 2021).

### Bacterial pathway, secretion, and taxonomy annotation

For the purposes of annotation, we selected the representative bacterial sequence of each cluster. If there was no bacterial representative, we sorted sequences by their status in Uniprot (reviewed/unreviewed) and by their length and chose the top sequence. Bacterial taxonomy information is associated with each UniRef90 cluster by HUMANN3 (Beghini et al., 2021). We submitted all bacterial protein sequences to the KofamKOALA (Aramaki et al., 2019) KEGG orthology search resource to obtain orthology and pathway annotations. To obtain secretion information, we used several sources: we submitted our bacterial sequences to EffectiveDB (Eichinger et al., 2016) in order to obtain predictions for EffectiveT3 (type 3 secretion based on signal peptide) and T4SEpre (type 4 secretion based on amino acid composition at the C-terminus). We used the single default cutoffs for T4SEpre, and chose the ‘selective’ (0.9999) cutoff for EffectiveT3. We obtained predictions for Sec and Tat pathway secretion using SignalP 5.0 (Almagro Armenteros et al., 2019) for Gram positive and Gram negative bacteria using default settings. Transmembrane proteins or signal peptides were predicted using TMHMM (Krogh et al., 2001) (v.2.0c), with a threshold of 19 or more expected number of amino acids in transmembrane helices. Localization to the cell wall was predicted using PSORTb 3.0 (Yu et al., 2010) with default settings. We annotated secretion systems in species associated with each bacterial cluster by examining the core or minimal components of each secretion system, by searching their genomes using KEGG orthologous groups for each system using string cutoffs (identity > 40%; e-value < 0.00001; coverage > 80%): T3SS: sctR (K03226), sctS (K03227), sctT (K03228), sctU (K03229), and sctV (K03230); T4SS: virB4 (K03199) and virD4 (K03205); Sec: secY (K03076), secE (K03073), and secG (K03075); and Tat: tatA (K03116) and tatC (K03118). We defined genomes in which have all minimal components of each system as organisms with functional corresponding secretion systems.

### Structural data for these microbiome-human PPIs

We measured the extent to which structural interfaces could be used to infer microbiome-human protein-protein interaction by using DIAMOND (Buchfink et al., 2015) to query all amino acid sequences submitted to PDB (identity > 70%; coverage > 50%). In order to identify interface residues between each pair of chains in the cocrystal structures, we first use NACCESS (http://www.bioinf.manchester.ac.uk/naccess/) to calculate the solvent accessibility of each residue in each chain. Chains with an accessible surface area of 15 Å or more are considered surface residues. We then calculate the change in accessible surface area for each residue when other chains in the same crystal structures are introduced. Residues which have a change in solvent accessible surface area above 1 Å are determined to be interface residues. Cases in which human protein and bacterial proteins match their respective chains exclusively are in Table S7. We highlight one example in which there are uniquely mapped chains, where 1p0s chains H and E match human coagulation factor X and bacterial Ecotin, respectively (Fig. S11).

To assess conservation of interface residues across bacterial members of the same UniRef cluster, we downloaded a list of all PDB structures which contain both human proteins and bacterial proteins, the UniRef50 cluster identifier for the bacterial protein, and all protein sequences in the corresponding cluster that also originate from bacterial proteomes from Uniprot. Using Clustal Omega, we then generated multiple sequence alignments for all the members of each UniRef50 clusters. We calculated interface residues on all pairs of chains in each structures and measured the BLOSUM62 similarity between bacterial interface residues and their corresponding amino acids in their respective UniRef50 cluster MSA. We then calculated the Jensen-Shannon divergence on the columns of the MSA containing interface residues.

## Supplemental Note 1

### Terminology used for gene-disease associations

The following terms from DisGeNet were used for each of the following broader disease annotations. For diabetes, we included all subtypes and diabetes-related phenotypes.

#### CRC

‘Colorectal Carcinoma’, ‘Colorectal Neoplasms’, ‘Adenocarcinoma of large intestine’, ‘Malignant tumor of colon’, ‘Hereditary Nonpolyposis Colorectal Neoplasms’, ‘Hereditary non-polyposis colorectal cancer syndrome’, ‘Hereditary Nonpolyposis Colorectal Cancer’, ‘Colorectal cancer, hereditary nonpolyposis, type 1’, ‘Hereditary nonpolyposis colorectal carcinoma’, ‘Colon Carcinoma’, ‘Colorectal Cancer, Susceptibility to, 4’, ‘Colorectal Cancer, Susceptibility to, on Chromosome 15’, ‘Colorectal Cancer, Hereditary Nonpolyposis, type 7 (disorder)’, ‘Colorectal Cancer, Hereditary Nonpolyposis, type 5’, ‘Colorectal Cancer, Hereditary Nonpolyposis, type 8’, ‘Colorectal Adenomatous Polyposis, Autosomal Recessive’, ‘Colorectal Cancer, Hereditary Nonpolyposis, type 4’, ‘Colorectal Cancer, Susceptibility to, 10’, ‘Colorectal Cancer, Susceptibility to, 12’, ‘Familial Colorectal Cancer Type X’, ‘Colorectal Cancer, Hereditary Nonpolyposis, type 6’, ‘Colorectal Cancer, Susceptibility to, 1’, ‘Oligodontia-Colorectal Cancer Syndrome’

#### Diabetes

‘Diabetes Mellitus, Experimental’, ‘Diabetic Nephropathy’, ‘Diabetes Mellitus, Non-Insulin-Dependent’, ‘Diabetes Mellitus, Insulin-Dependent’, ‘Diabetes, Autoimmune’, ‘Brittle diabetes’, ‘Diabetes Mellitus, Ketosis-Prone’, ‘Diabetes Mellitus, Sudden-Onset’, ‘Diabetic Retinopathy’, ‘Diabetic Cardiomyopathies’, ‘Diabetic cystopathy’, ‘Diabetes Mellitus’, ‘Complications of Diabetes Mellitus’, ‘Neonatal diabetes mellitus’, ‘Gestational Diabetes’, ‘Alloxan Diabetes’, ‘Streptozotocin Diabetes’, ‘Prediabetes syndrome’, ‘Diabetic Angiopathies’, ‘Microangiopathy, Diabetic’, ‘Diabetes Mellitus, Noninsulin-dependent, 1 (disorder)’, ‘Diabetic Neuropathies’, ‘Symmetric Diabetic Proximal Motor Neuropathy’, ‘Asymmetric Diabetic Proximal Motor Neuropathy’, ‘Diabetic Mononeuropathy’, ‘Diabetic Polyneuropathies’, ‘Diabetic Amyotrophy’, ‘Diabetic Autonomic Neuropathy’, ‘Diabetic Asymmetric Polyneuropathy’, ‘Diabetic Neuralgia’, ‘Nephrogenic Diabetes Insipidus’, ‘Diabetes Mellitus, Insulin-Dependent, 22 (disorder)’, ‘Microcephaly, Epilepsy, and Diabetes Syndrome’, ‘Diabetes’, ‘Diabetes Mellitus, Insulin-Dependent, 12’, ‘Microvascular Complications of Diabetes, Susceptibility to, 3 (finding)’, ‘Diabetes Mellitus, Neonatal, with Congenital Hypothyroidism’, ‘Phosphate Diabetes’, ‘Diabetic encephalopathy’, ‘Microvascular Complications of Diabetes, Susceptibility to, 2 (finding)’, ‘Insulin-resistant diabetes mellitus’, ‘Lymphedema-Distichiasis Syndrome with Renal Disease and Diabetes Mellitus’, ‘Lipoatrophic Diabetes Mellitus’, ‘Pregnancy in Diabetics’, ‘Maturity onset diabetes mellitus in young’, ‘Maturity-Onset Diabetes of the Young, type 14’, ‘Latent Autoimmune Diabetes in Adults’, ‘Monogenic diabetes’, ‘Diabetes mellitus autosomal dominant type II (disorder)’, ‘ Diabetes Mellitus, Permanent Neonatal’, ‘Diabetes Insipidus’, Microvascular Complications of OF Diabetes, Susceptibility to, 7 (finding)’, ‘Renal cysts and diabetes syndrome’, ‘Maturity-Onset Diabetes of the Young, Type 1’, ‘Fanconi Renotubular Syndrome 4 with Maturity-onset Diabetes of the Young’, ‘Transient neonatal diabetes mellitus’, ‘Diabetes Mellitus, Transient Neonatal, 1’, ‘Diabetes Mellitus, Insulin-Dependent, 2’, ‘diabetes (mellitus) due to autoimmune process’, ‘Diabetes (mellitus) due to immune mediated pancreatic islet beta-cell destruction’, ‘Idiopathic Diabetes (Mellitus)’, Microvascular Complications of Diabetes, Susceptibility to, 4 (finding)’, ‘Diabetes Mellitus, Insulin-Dependent, 10’, ‘Acquired Nephrogenic Diabetes Insipidus’, ‘Congenital Nephrogenic Diabetes Insipidus’, ‘Nephrogenic Diabetes Insipidus, Type I’, ‘Nephrogenic Diabetes Insipidus, Type II’, ‘ADH-Resistant Diabetes Insipidus’, ‘Diabetic Ketoacidosis’, ‘Non-insulin-dependent diabetes mellitus with unspecified complications’, ‘Diabetes Mellitus, Permanent Neonatal, with Neurologic Features’, ‘Developmental Delay, Epilepsy, and Neonatal Diabetes’, ‘Maturity-onset diabetes of the young, type 10’, ‘Diabetes Mellitus, Insulin-Resistant, with Acanthosis Nigricans’, ‘Maturity-onset Diabetes of the Young, type IV (disorder)’, ‘Diabetes Mellitus, Transient Neonatal, 3 (disorder)’, ‘Maturity-onset Diabetes of the Young, type 13’, ‘Diabetes Mellitus, Insulin-Dependent, 5’, ‘ Diabetes Mellitus, Insulin-Dependent, 7’, ‘Maturity-onset Diabetes of the Young, type 6 (disorder)’, ‘Gastroparesis with diabetes mellitus’, ‘Other specified diabetes mellitus with unspecified complications’, ‘Insulin-dependent diabetes mellitus secretory diarrhea syndrome’, ‘Severe nonproliferative diabetic retinopathy’, ‘Microvascular Complications of Diabetes, Susceptibility to, 5 (finding)’, ‘Central Diabetes Insipidus’, ‘Ataxia, Combined Cerebellar and Peripheral, with Hearing Loss and Diabetes Mellitus’, ‘Maturity-onset diabetes of the young, type 11’, ‘Microvascular Complications of Diabetes, Susceptibility to, 6 (finding)’, ‘Diabetes Mellitus, Transient Neonatal, 2 (disorder)’, ‘Maturity-onset Diabetes of the Young, type 3 (disorder)’, ‘Diabetes Mellitus, Insulin-Dependent, 20 (disorder)’, ‘Proliferative diabetic retinopathy’, ‘Microvascular Complications of Diabetes, Susceptibiltiy to, 1(finding)’, ‘Maturity-onset Diabetes of the Young, type type 7 (disorder)’, ‘Diabetes Mellitus, Noninsulin-dependent, 5’

#### Autoimmune

‘Autoimmune hemolytic anemia’, ‘Autoimmune Diseases’, ‘Autoimmune state’, ‘Celiac Disease’, ‘Lupus Erythematosus, Systemic’, ‘Diabetes, Autoimmune’, ‘Autoimmune Chronic Hepatitis’, ‘Rheumatoid Arthritis’, ‘Ankylosing spondylitis’, ‘Multiple Sclerosis’, ‘Autoimmune Lymphoproliferative Syndrome’, ‘Experimental Autoimmune Encephalomyelitis’, ‘Lupus Erythematosus, Cutaneous’, ‘Chilblain lupus 1’, ‘Multiple Sclerosis, Acute Fulminating’, ‘Autoimmune thyroiditis’, ‘Autoimmune Lymphoproliferative Syndrome Type 2B’, ‘Autoimmune Interstitial Lung, Joint, and Kidney Disease’, ‘Lupus Vulgaris’, ‘Lupus Erythematosus, Discoid’, ‘Lupus Erythematosus’, ‘Rheumatoid Arthritis, Systemic Juvenile’, ‘Neuritis, Autoimmune, Experimental’, Systemic Lupus Erythematosus 16’, ‘Ankylosing spondylitis and other inflammatory spondylopathies’, ‘Lupus Vasculitis, Central Nervous System’, ‘Lupus Meningoencephalitis’, ‘Neuropsychiatric Systemic Lupus Erythematosus’, ‘Lupus Nephritis’, ‘Vitiligo-associated Multiple Autoimmune Disease Susceptibility 1 (finding)’, Chilblain Lupus 2’, ‘Latent Autoimmune Diabetes in Adults’, ‘Vitiligo-associated Multiple Autoimmune Disease Susceptibility 6’, ‘Autoimmune Disease, Susceptibility to, 1’, ‘Autoimmune Hepatitis with Centrilobular Necrosis’, ‘Polyendocrinopathies, Autoimmune’, ‘Polyglandular Type I Autoimmune Syndrome’, ‘Autoimmune Syndrome Type II, Polyglandular’, ‘Polyglandular Type III Autoimmune Syndrome’, ‘Autoimmune Polyendocrinopathy Syndrome, Type I, Autosomal Dominant’, ‘Autoimmune Polyendocrinopathy Syndrome, type I, with Reversible Metaphyseal Dysplasia’, ‘Autoimmune polyendocrinopathy syndrome, type 1’, ‘Multiple Sclerosis, Acute Relapsing’, ‘Multiple Sclerosis, Relapsing-Remitting’, ‘diabetes (mellitus) due to autoimmune process’, ‘Autoimmune Lymphoproliferative Syndrome, Type IA’, ‘Ras-associatedAutoimmune Leukoproliferative Disorder’, ‘Autoimmune Lymphoproliferative Syndrome Type 1, Autosomal Dominant’, ‘Autoimmune Diseases of the Nervous System’, ‘Autoimmune Disease, Susceptibility to, 6’, ‘Autoimmune Lymphoproliferative Syndrome, Type III’, ‘Alpha/Beta T-cell Lymphopenia with Gama/Delta T-cell Expansion, Severe Cytomegalovirus Infection, and Autoimmunity, ‘Idiopathic Autoimmune Hemolytic Anemia’, ‘Autoimmune Disease, Multisystem, Infantile-onset, 1’, ‘Systemic Lupus Erythematosus, Multisystem, 11’, ‘T-cell Immunodeficiency, Recurrent Infections, and Autoimmunity with or without Cardiac Malformations’, ‘T-cell Immunodeficiency, Recurrent Infections, Autoimmunity, and Cardiac Malformations’, ‘Hyperthyroidism, Nonautoimmune’, ‘Autoimmune Disease, Multisystem, Infantile-onset, 2’, ‘Autoimmune Disease, Multisystem, with facial dysmorphism’, ‘Syndromic multisystem autoimmune disease due to itch deficiency’, ‘Autoimmune Lymphoproliferative Syndrome, Type IIA’, ‘Immunodeficiency, Common Variable, 8 with Autoimmunity’

#### Obesity

‘Obesity’, ‘Pediatric Obesity’, ‘Adolescent Obesity’, ‘Childhood Overweight’, ‘Infantile Obesity’, ‘Infant Overweight’, ‘Adolescent Overweight’, ‘Abdominal obesity metabolic syndrome’, ‘Obesity, Morbid’, ‘Obesity, Hyperphagia, and Developmental Delay’, ‘Obesity, Abdominal’, ‘Mental Retardation, Epilectic Seizures, Hypogonadism and Hypogenitalism, Microcephaly, and Obesity (disorder)’, ‘Obesity, Susceptibility to’, ‘Obesity, Visceral’, ‘Overweight’, ‘Obesity due to melanocortin 4 receptor deficiency’, ‘ABDOMINAL Obesity-Metabolic Syndrome 1’, ‘Developmental Delay, Intellectual Disability, Obesity, and Feautres’, ‘Spastic Paraplegia, Intellectual disability, nystagmus, and Obesity’, ‘Retinal Dystrophy and Obesity ‘, ‘Childhood-onset truncal obesity’, ‘Morbid Obesity and Spermatogenic Failure’, ‘Abdominal Obesity-Metabolic Syndrome 3’

#### IBD

‘Ulcerative Colitis’, ‘Crohn Disease’, ‘Colitis’, "Crohn’s disease of large bowel", ‘Inflammatory Bowel Diseases’, ‘Necrotizing Enterocolitis’, "Crohn’s disease of the ileum", ‘IIeocolitis’, ‘Inflammatory Bowel Disease 17’, ‘Chronic left-sided ulcerative colitis’, ‘Inflammatory Bowel Disease 12’, ‘Inflammatory Bowel Disease 19’, ‘Enterocolitis’, ‘Enterocolitis, Neutropenic’, ‘Inflammatory bowel disease 28, Autosomal Recessive’, ‘Inflammatory bowel disease 25, autosomal recessive’, ‘Inflammatory Bowel Disease 14’, ‘Inflammatory Bowel Disease 13’, ‘Inflammatory Bowel Disease 10’,’Inflammatory Bowel Disease 29’, ‘Autoinflammation with Infantile Enterocolitis’ ‘Crohn Disease-associated Growth Failure, Susceptibility to (finding)’, ‘Neutropenic colitis’,;Inflammatory Bowel Disease, Immunodeficiency, and encephalopathy’, ‘Inflammatory Bowel Disease, Immunodeficiency, and Ecnephalopathy’, ‘Inflammatory Bowel Disease 16’

## Supplementary Figures

**Figure S1.**
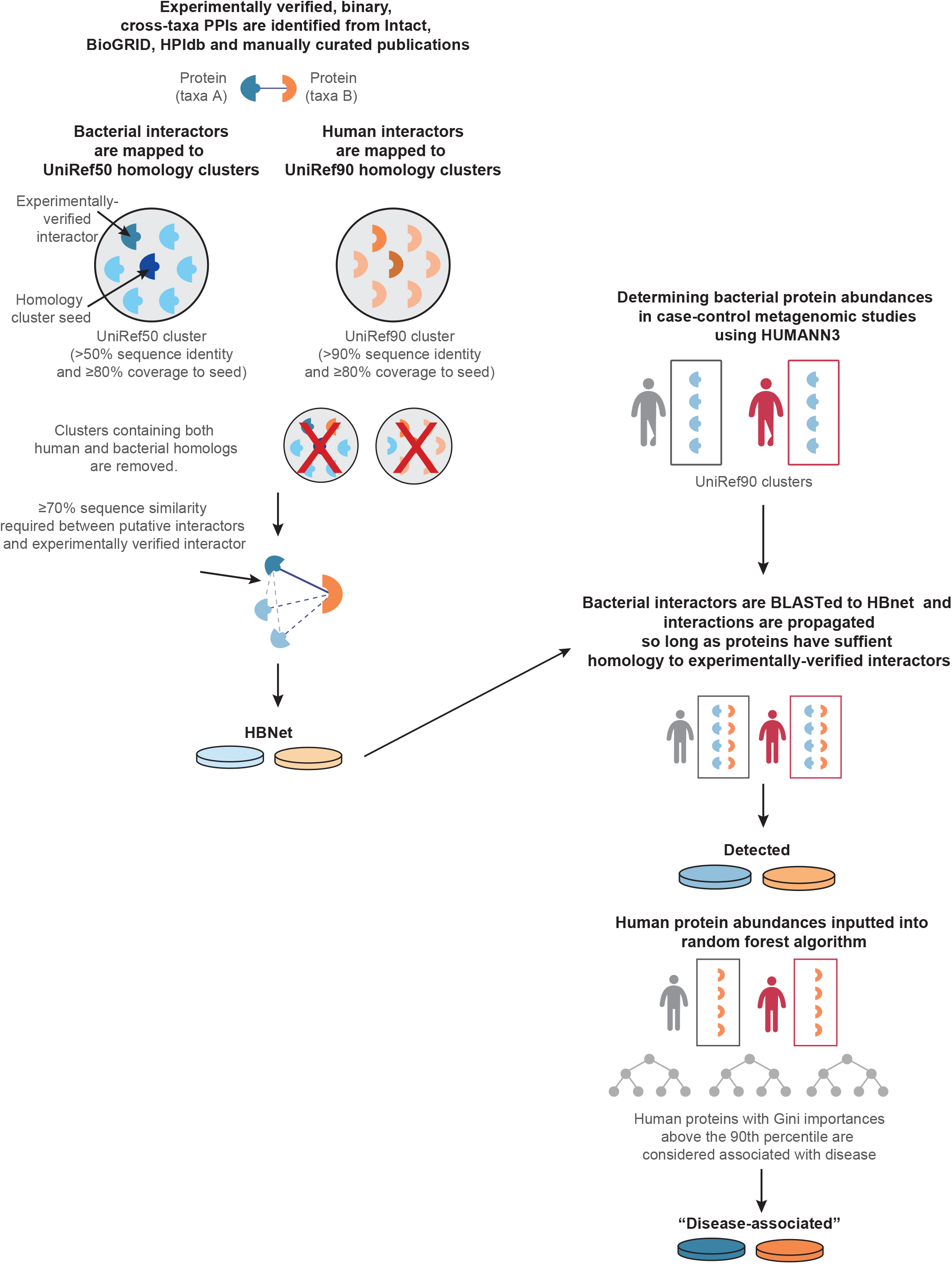
Few bacterial-human interaction sequences populate the Protein Data Bank. A Venn diagram describing the number of detected bacterial clusters and human interactors in the nine metagenomic cohorts that have any matching structure (using BLASTp) in the PDB to at least one chain (medium blue) and whether their homologous structures appear on the same PDB cocrystal structure (dark blue). Only one PDB structure showed non-overlapping homology to both a human and bacterial protein.

**Figure S2.**
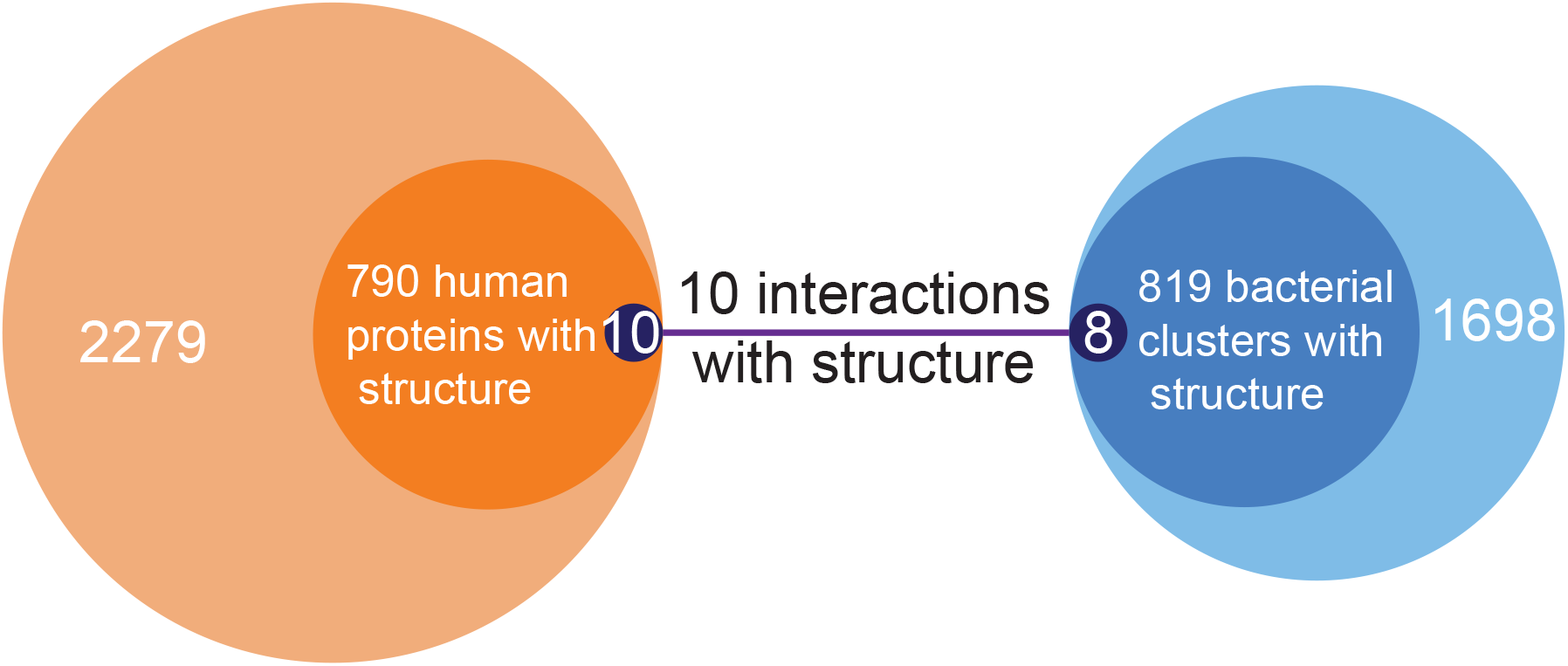
An outline of our homology mapping procedure and alignment. Depiction of the interaction network inference and protein detection pipeline for bacterial/microbiome (blue)-human (orange) PPIs.

**Figure S3.**
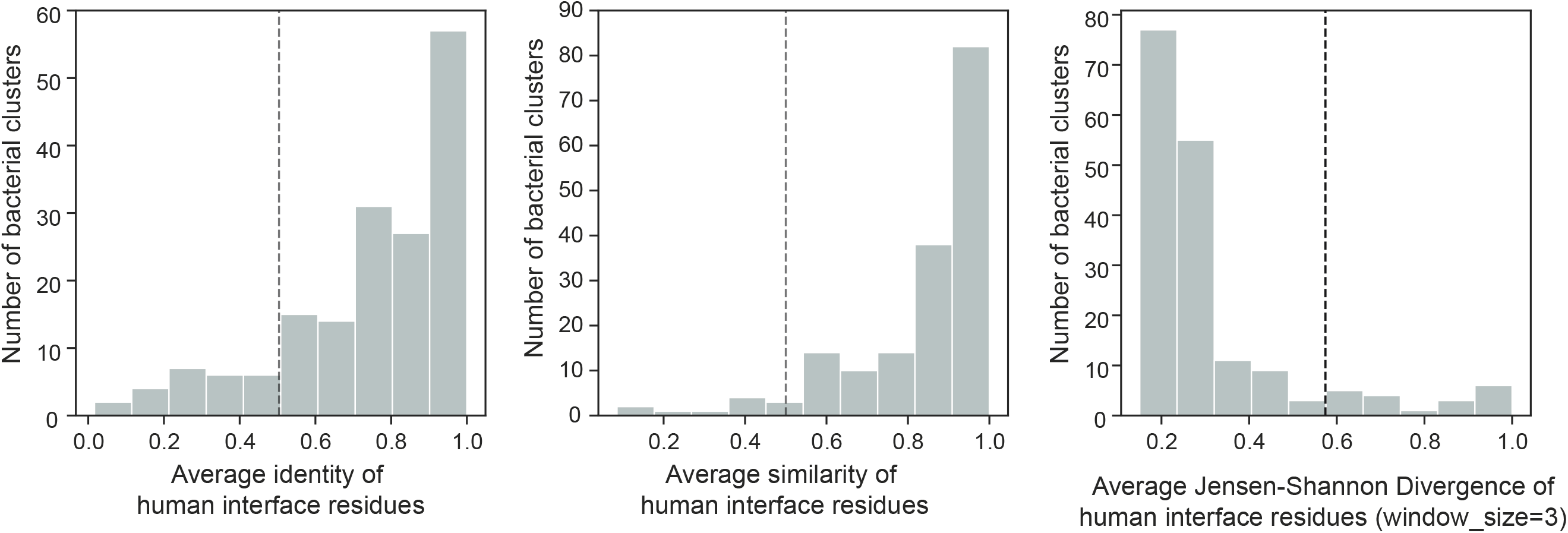
Interface similarity between bacterial proteins within a UniRef cluster. Similarity, identity, and Jensen-Shannon divergence of interface residues across all bacterial members of the same UniRef cluster sourced from all cocrystal structures in the PDB with human and bacterial interactors.

**Figure S4.**
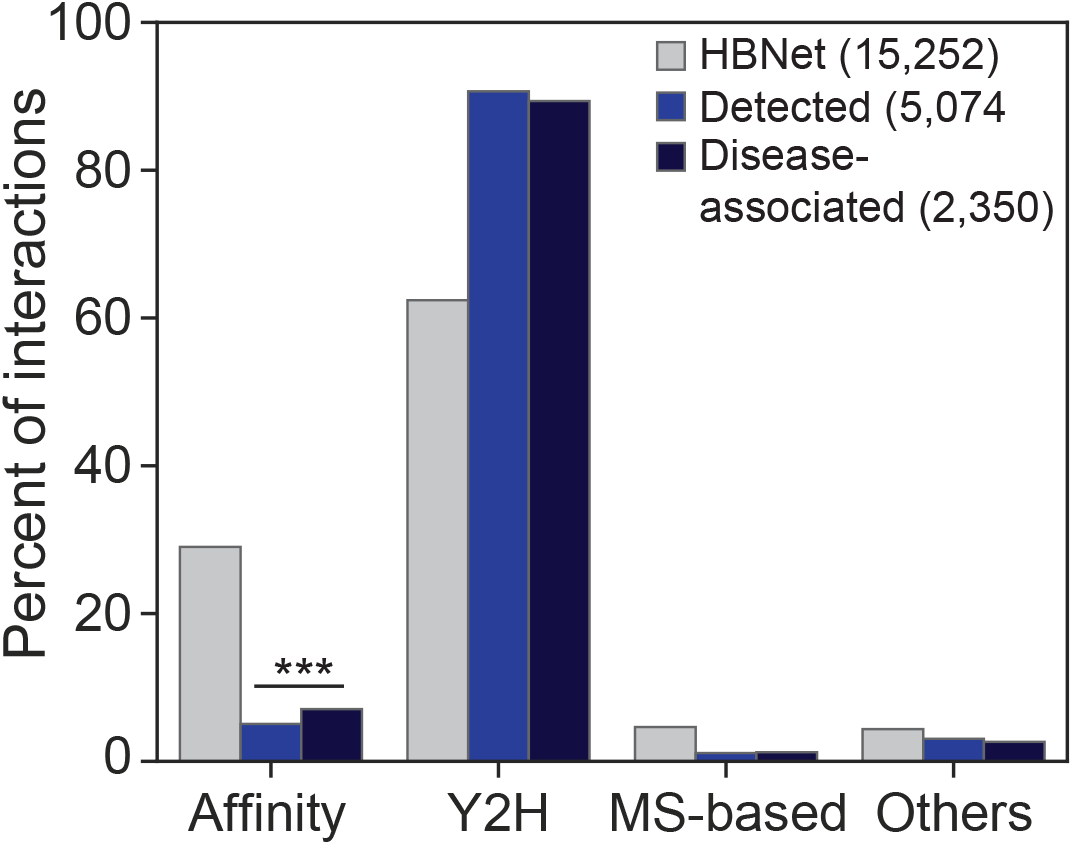
Disease-associated interactions are enriched for those based on affinity-based methods. The three largest categories of detection methods are shown (affinity-based methods, yeast-2-hybrid, mass spectrometry methods) as well as ‘Other’. p-values are only shown between ‘Detected’ and ‘Disease-associated’ and are depicted by: * p<0.05; ** p<0.01; *** p<0.001; **** p<0.0001 (Chi-square test). Total numbers of each set are noted in the legend.

**Figure S5.**
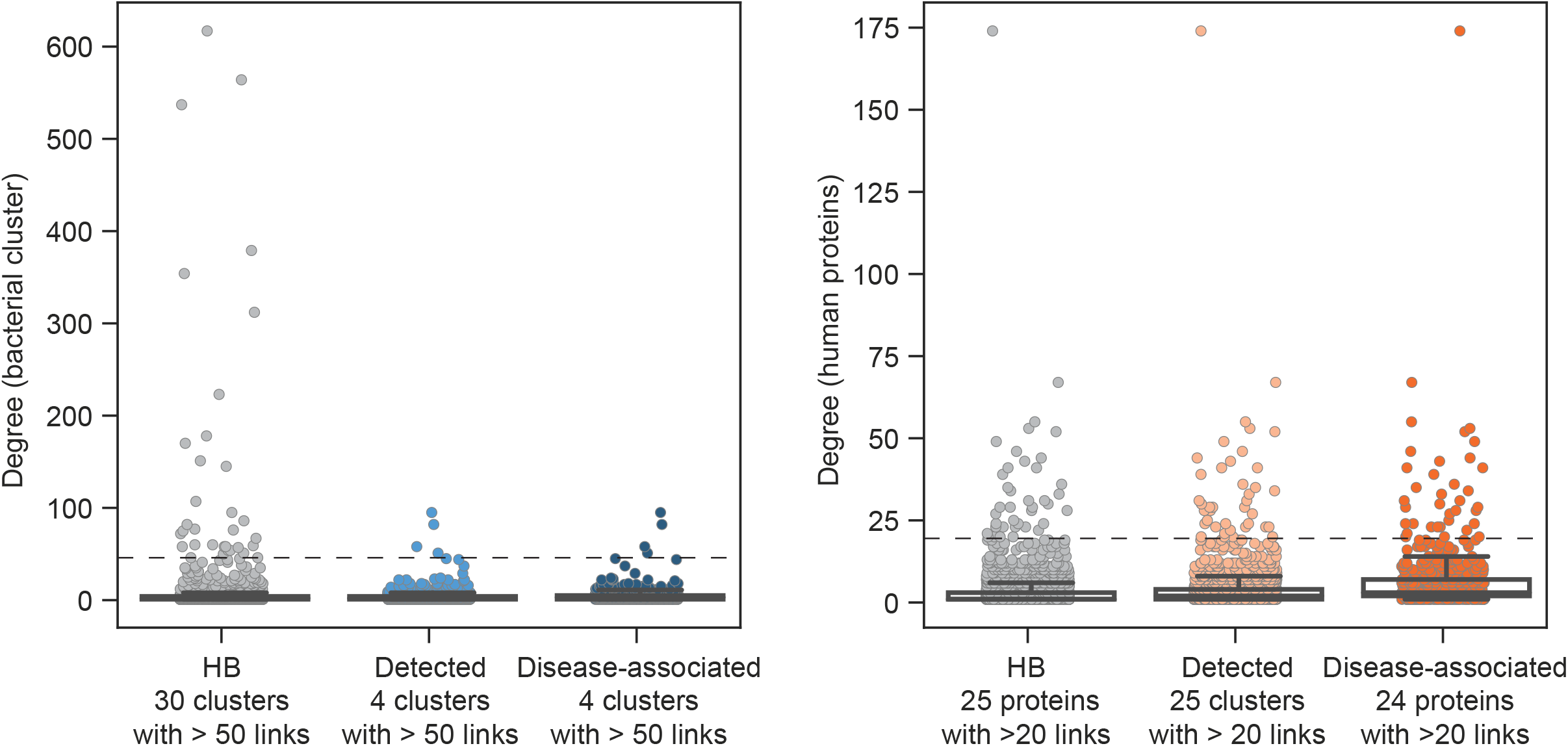
Degree distribution for bacterial protein clusters and human proteins. The degree distribution per bacterial protein cluster (left) or human protein (right) in the HBNet, Detected or Disease-associated subsets.

**Figure S6.**
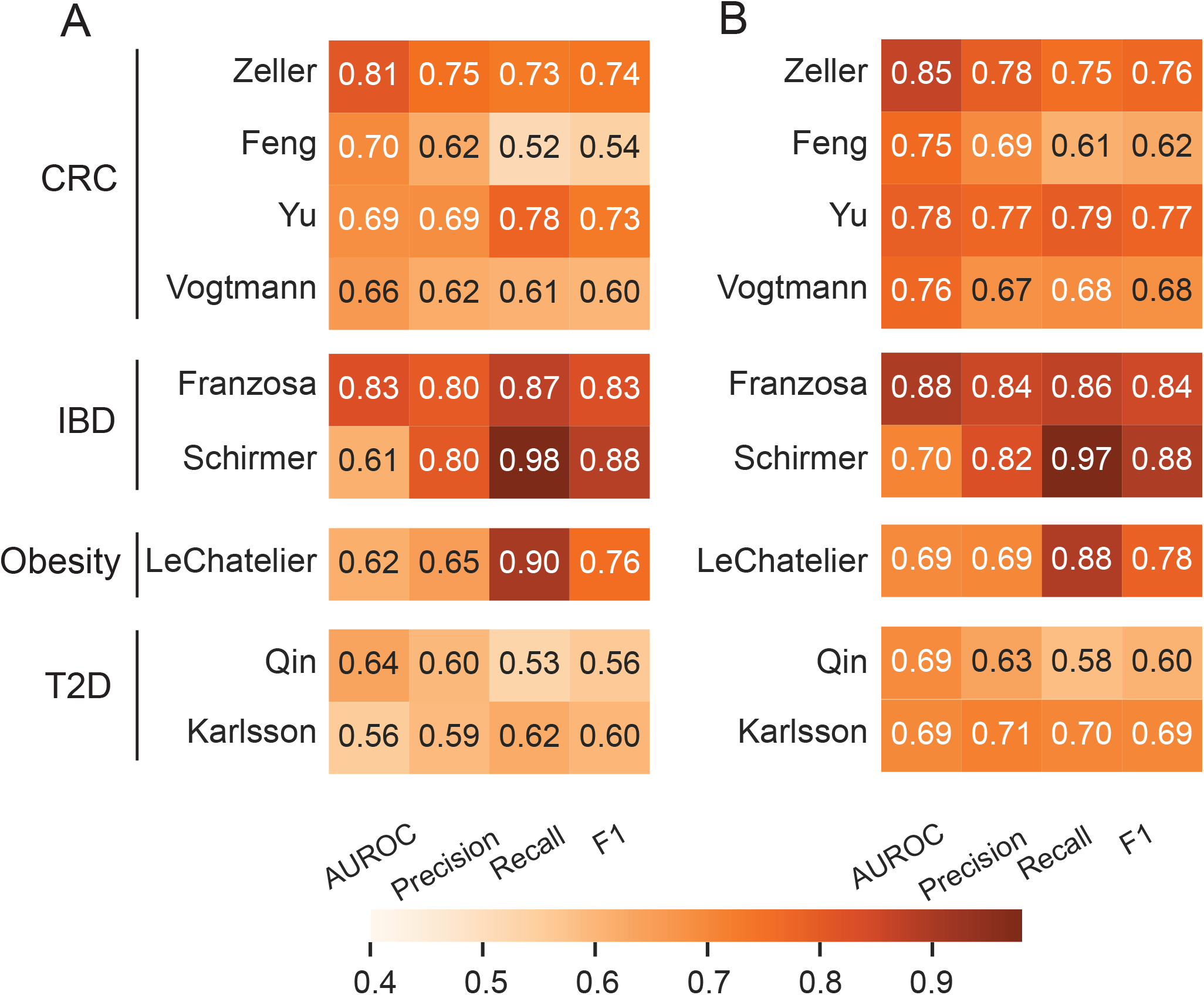
Performance metrics of the random forest (RF) classifier. (A) A heatmap of area under the receiver operating characteristic curve (AUROC), precision, recall, and F1-scores for random forests on the putative human interactors with the microbiomes of each metagenomic study with grid search-based hyper-parameter tuning, evaluated using five-fold cross validation. (B) Performance metrics of the RF classifier using only features above the 90^th^ percentile.

**Figure S7.**
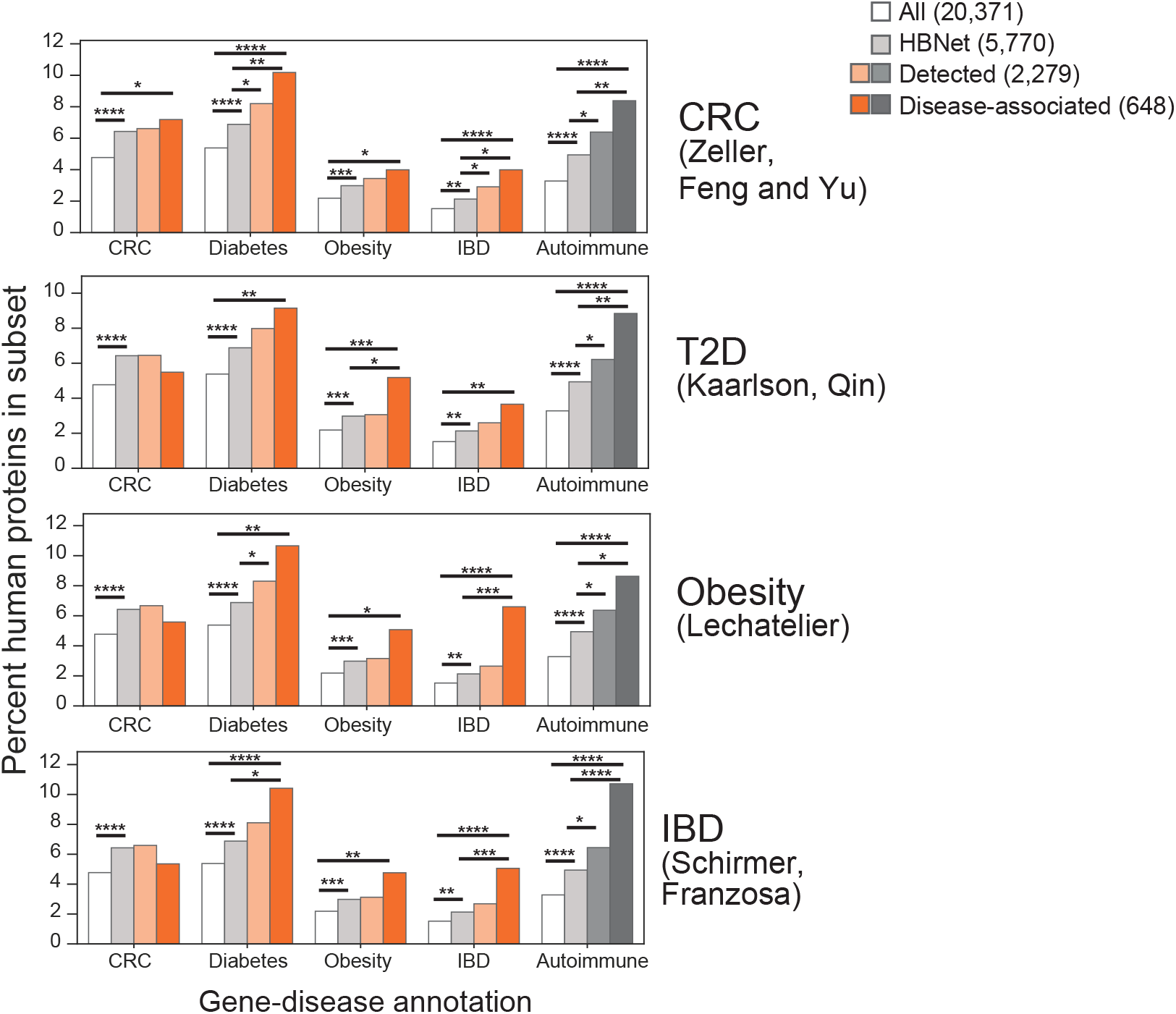
Gene-disease annotations are specific to each disease cohort. (A) The proportions of human proteins implicated in disease, according to their GDAs in DisGeNET (only GDAs with scores over 0.1 were considered) and grouped according to disease-specific cohorts, in the following subsets: all reviewed human proteins (totaling 20,371 proteins); HBNet (5,770 proteins); human interactors with detected bacterial proteins (2,279 proteins); and those human interactors with feature importances above the 90th percentile in their respective cohorts (648 unique proteins). p-values are depicted by: * p<0.05; ** p<0.01; *** p<0.001; **** p<0.0001 (Chi-square test). Total numbers of each set are noted in the legend.

**Figure S8.**
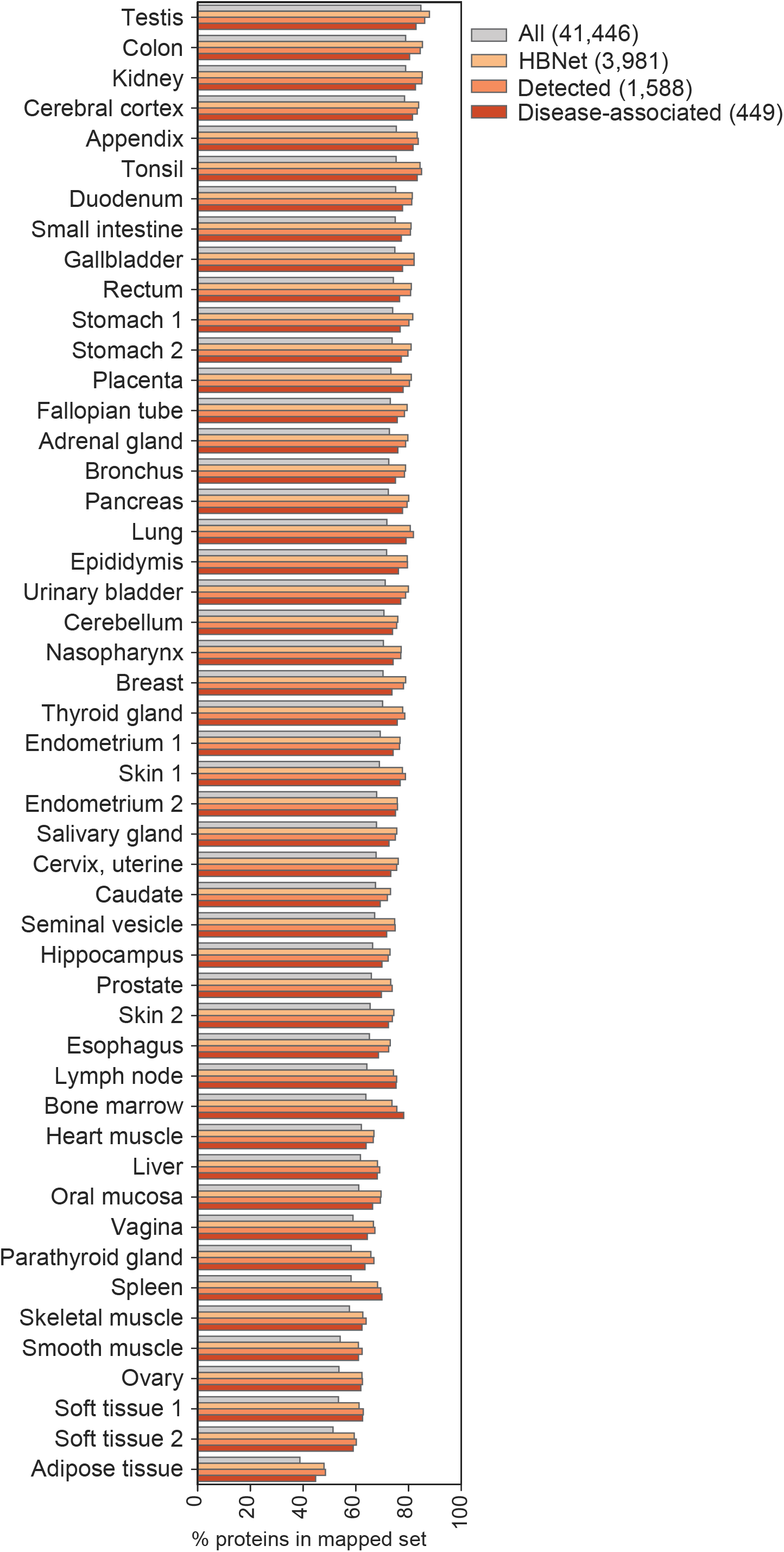
Protein localization and protein expression according to human tissue. Protein localization according to tissue, as annotated by the Human Protein Atlas. Only those with “enhanced”, “supported” or “approved” annotations were included. Total numbers of each set are noted in the legend.

**Figure S9.**
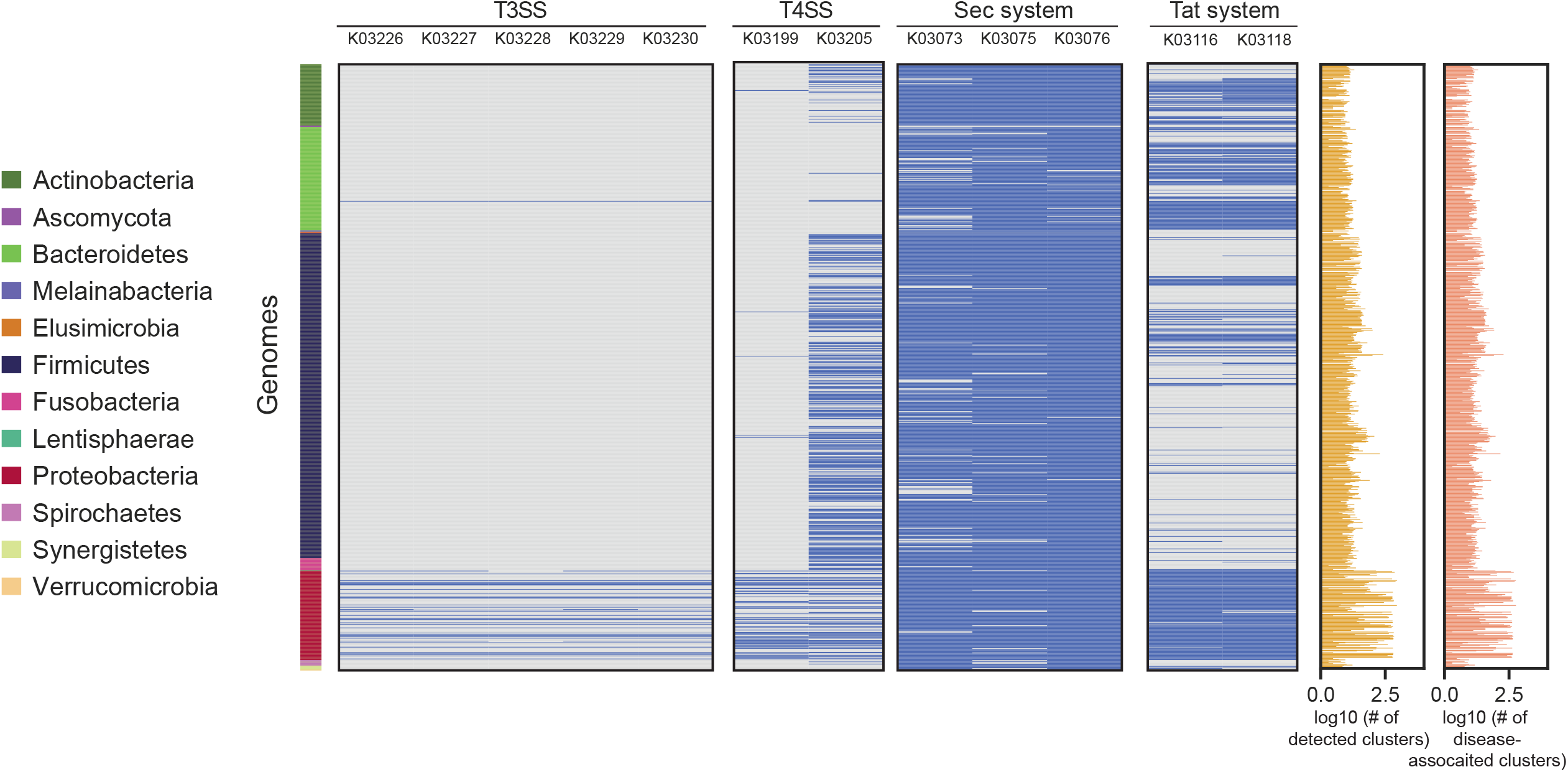
Secretion systems distribution varies across bacterial species. A heatmap (present/absent) of the required components for each secretion system (denoted using their KO numbers) present in each bacterial species (colored by phylum to the left) with at least one detected protein associated with bacterial protein clusters in nine case-control cohort studies. The actual number of detected and disease-associated protein cluster representatives for each bacteria in any of the nine metagenomic studies is plotted to the right.

**Figure S10.**
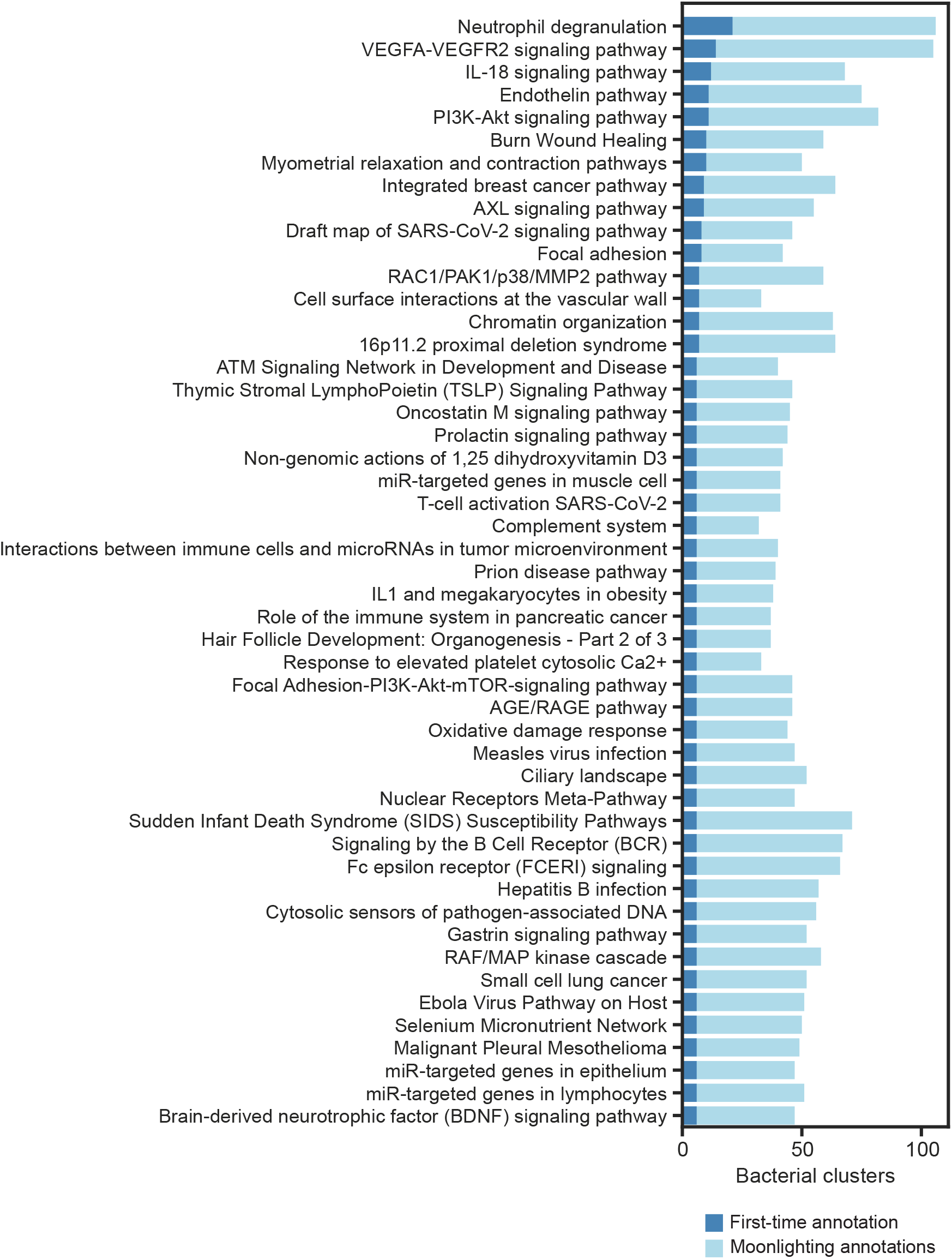
Bacterial clusters gain putative human-relevant functions. Human pathways (annotated using WikiPathways) significantly enriched (FDR-adjusted p-values < 0.05) in either HBNet, the human proteins targeted by bacterial clusters detected in the metagenomic studies, or those human targets associated with disease in the metagenomic case-control cohort studies (disease-associated). 953 out of 1,102 metagenomic cohort-associated human proteins were able to be annotated. Note that each bacterial protein cluster may gain multiple annotations, according to the roles of their human interactor(s).

**Figure S11.**
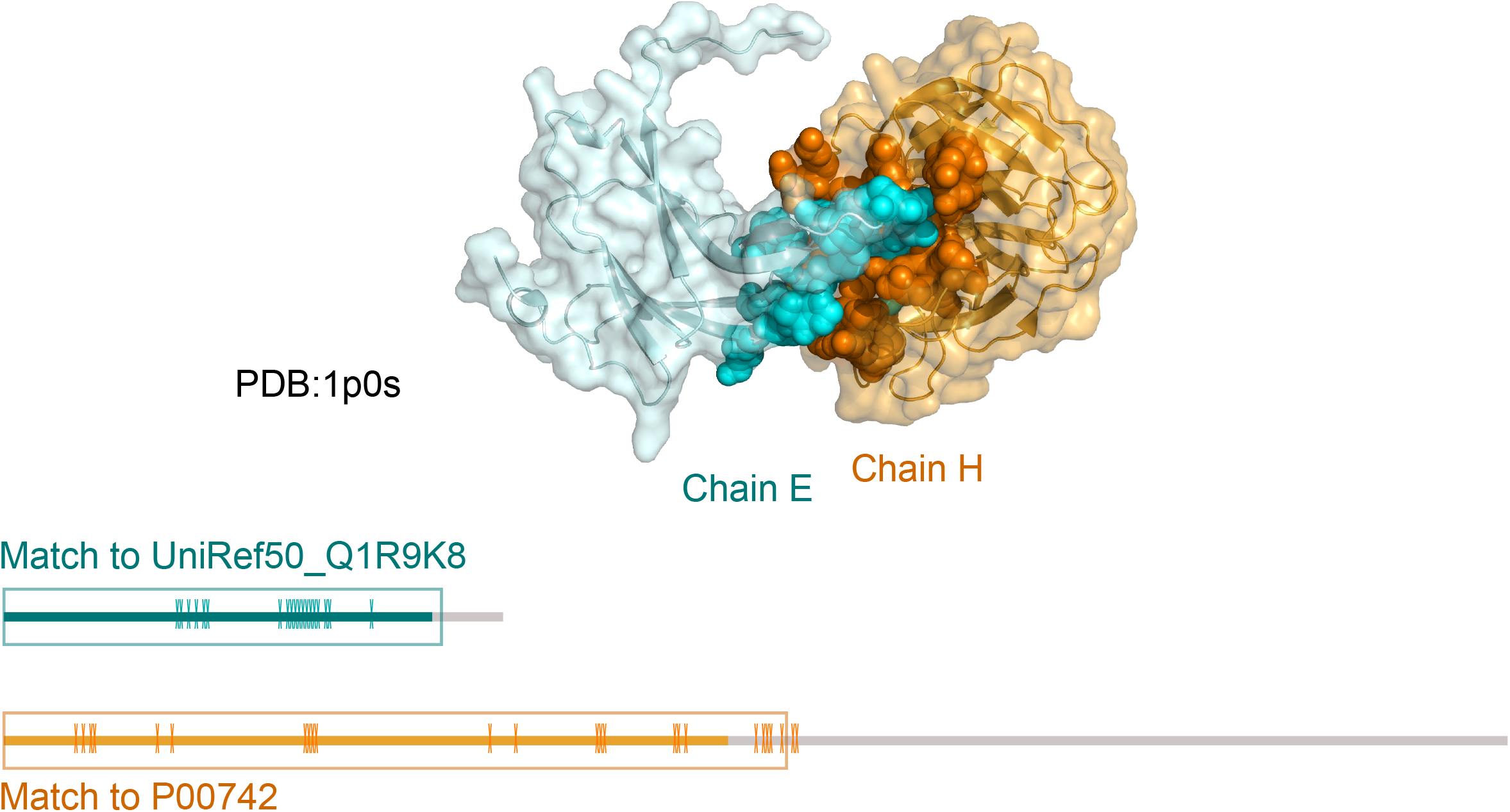
Cocrystal structure of blood coagulation factor Xa in complex with Ecotin M84R. Cluster Uniref50_Q1R9K8 contains several bacterial ecotins detected in human metagenomes. Using BLAST, we found high-quality matches between members of this cluster and the structure 1p0s:E (Ecotin precursor M84R) in the PDB (identity of 97.2%, eval=10-75). Our putative interactor to this cluster, coagulation factor X (P00742) likewise matched structure 1p0s:H (coagulation factor X precursor) (identity of 100%, eval=3.8×10-150). Chain E is shown in blue, and chain H in orange, with their interface residues highlighted as spheres. The linear model of both proteins is shown underneath. The linear model’s colored areas indicate the part of the proteins that were crystallized in this PDB, while the greyed-out areas indicate non-crystallized spans. The squares indicate the range of the BLAST match between our query proteins and the PDB reference sequences. Finally, ticks on the linear model indicate the location of interface residues as detected in this model. There are currently not enough published structures to perform this analysis on all interactions involving detected bacterial genes (Fig. S2, Table S7).

## Supplementary Tables

**Table S1. Extended information on known experimentally verified host-microbiome interactions with evidence for a role in cellular physiology and/or human health.** Information on the interaction detection method for human-microbiome PPIs that have been shown to affect cell physiology and/or human health.

**Table S2. Metagenomic samples used in this research.** For each study, we list the sample numbers and labels in the cohort study.

**Table S3. Disease-associated human-microbiome PPIs.** Human-microbiome PPIs are listed according to their UniProt and UniRef50 identifiers, human and bacterial protein names.

**Table S4. Number of human interactors according to the source of the experimentally-verified interactors.** The number of human interactors, according to the species sourcing the initial experimentally verified interacting protein.

**Table S5. Human interactors that are known drug targets.** For each disease-associated human protein, we list the drug interactor (annotated using DrugCentral and DrugBank) and the study in which it was found to be important.

**Table S6. Extended information for bacterial proteins targeting known drug targets in Figure 4.** Bacterial clusters depicted in Fig. 4 are listed with their UniRef number and detected taxa, according to HUMANN3.

**Table S7. Cocrystal structures representing interactions in our dataset.** All pairs of detected bacterial proteins and human proteins in the nine metagenomic datasets that have BLASTp matches to two different chains within the same PDB cocrystal structure (totaling 8 bacterial protein clusters and 10 human proteins). This list includes structures with at least one chain exclusive to each bacterial and human proteins.

